# rec-Y2H matrix screening reveals a vast potential for direct protein-protein interactions among RNA binding proteins

**DOI:** 10.1101/2020.09.14.296160

**Authors:** Benjamin Lang, Jae-Seong Yang, Mireia Garriga-Canut, Silvia Speroni, Maria Gili, Tobias Hoffmann, Gian Gaetano Tartaglia, Sebastian P. Maurer

## Abstract

RNA-binding proteins (RBPs) are crucial factors of post-transcriptional gene regulation and their modes of action are intensely investigated. At the center of attention are RNA motifs that guide where RBPs bind. However, sequence motifs are often poor predictors of RBP-RNA interactions *in vivo*. It is hence believed that many RBPs recognize RNAs as complexes, to increase specificity and regulatory possibilities. To probe the potential for complex formation among RBPs, we assembled a library of 978 mammalian RBPs and used rec-Y2H screening to detect direct interactions between RBPs, sampling > 600 K interactions. We discovered 1994 new interactions and demonstrate that interacting RBPs bind RNAs adjacently *in vivo*. We further find that the mRNA binding region and motif preferences of RBPs can deviate, depending on their adjacently binding interaction partners. Finally, we reveal novel RBP interaction networks among major RNA processing steps and show that splicing impairing RBP mutations observed in cancer rewire spliceosomal interaction networks.

**Graphical abstract:** 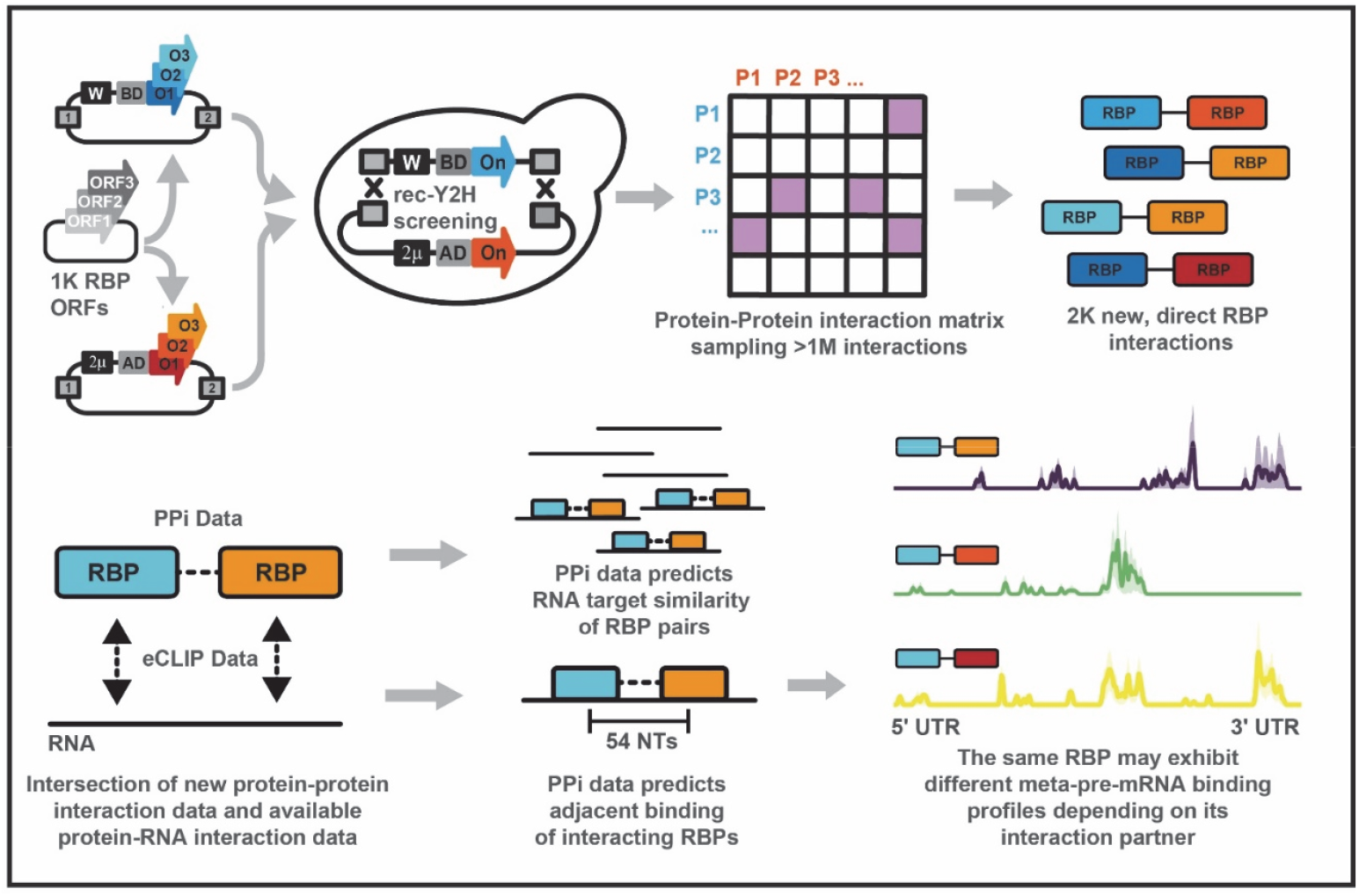

## Introduction

RNA-binding proteins (RBPs) interact with RNAs from the first moments of transcription and guide RNA processing, nuclear export, cytoplasmic localization, translation and decay. As such, RBPs are crucial for guiding gene expression in time, space and depending on the cellular context. Hundreds of RBPs have been discovered so far (Hentze et al., 2018) and numerous methods have been developed to understand which RNAs are bound by which RBPs and where (Garriga-Canut et al., 2019; König et al., 2010; Lambert et al., 2014; Van Nostrand et al., 2016; Ray et al., 2009). The locations where RBPs bind an RNA are guided by primary RNA sequences (Ray et al., 2013), tertiary structures (Stefl et al., 2010; Tan et al., 2013) and post translational modifications (Zaccara et al., 2019). These locations can provide information about the function of RBPs (Mukherjee et al., 2019b). Still, understanding the determinants of RBP-RNA interactions is a controversial and intense field of research. Predicting where an RBP might bind by using RNA sequences, that are often degenerated, is not straight forward; many RBP motifs exist on a large fraction of the transcriptome, but only a small percentage of these motifs are actually bound *in vivo* (Li et al., 2010). Possible explanations for this observation are that an RBP motif can be sequestered in inaccessible secondary structures (Taliaferro et al., 2016), that motifs are sterically blocked by other RBPs that bind an adjacent motif with higher affinity (Zarnack et al., 2013), or that a larger sequence context is important for RBP target selection (Dominguez et al., 2018; Hiller et al., 2006; Li et al., 2010). Another possibility is that many RBPs act in complexes (Achsel and Bagni, 2016; Quattrone and Dassi, 2019; Sternburg and Karginov, 2020) and that only when motifs for two or more RBPs are present in the right spacing and orientation, efficient binding can occur. Complex formation can increase specificity, affinity and avidity, thereby increasing the precision of regulation (Singh and Valcárcel, 2005). Also, combinatorial binding of RBPs increases the possibilities for regulatory complexity and enables encoding of multiple checkpoints (only when two or more RBPs bind, a process is triggered). Such combinatorial sequence recognition is often observed for DNA-binding transcription factors, which possess a range of cooperative DNA binding modes (Morgunova and Taipale, 2017; Stampfel et al., 2015) and can even alter their motif preference depending on their interaction partner (Cirillo et al., 2015; Jolma et al., 2015). In the case of RBPs, a few examples exist illustrating how RBP-RBP interactions trigger high-affinity binding, enable a specific process or alter the function of an RBP depending on its interaction partner (Achsel and Bagni, 2016; Fernandez et al., 2015; Hennig et al., 2014; Müller-McNicoll et al., 2016; Piqué et al., 2008). To which extent combinatorial RNA-binding mechanisms are employed by RBPs, however, is not systematically understood. A major limitation is an RBPome-scale understanding of which RBPs directly interact. Detecting direct RBP interactions has been difficult, potentially due to the often disordered-domain mediated, transient interactions amongst them, which are currently highlighted by the intense research on liquid condensates formed by RBPs (Järvelin et al., 2016). The problem is further illustrated by a number of affinity purification-mass spectrometry (AP-MS) studies. Here, the removal of RNA led to the loss of many interacting proteins (Brannan et al., 2016; Fritzsche et al., 2013; Kanai et al., 2004; Mallardo et al., 2003), possibly due to the washing steps required which may lead to loosing of weakly and transiently interacting RBPs. A major advance are recently improved *in vivo* proximity biotinylation techniques (Fazal et al., 2019; Mukherjee et al., 2019a). However, these techniques cannot distinguish between direct and indirect interactions among RBPs either.

To systematically search for direct RBP interactions, we hence used high-throughput recombination Yeast two-hybrid (rec-Y2H) matrix screening (Yang et al., 2018) to probe >1 million interactions between 978 mammalian RNA-binding proteins. As the spatial organization of the transcriptome is emerging as an important aspect of gene regulation, we added 76 microtubule-associated proteins (MAPs), i.e. motor proteins and cargo adaptors to the screen library to put extra emphasis on detecting new interactions between RBPs and the microtubule-based transport machinery. Using an improved rec-Y2H analysis pipeline, which increases sensitivity while keeping the same specificity (see Methods), we detected 1,994 novel interactions and validated 422 previously reported interactions. We show that binary RBP interactions detected by our screen can predict co-binding of RNAs and even proximal binding of RBPs at transcriptome-scale, likely revealing new functional RBP complexes. RBPs that show significant binding in immediate proximity relative to randomized samples often engage in multiple interactions that can alter their pre-mRNA region preference, which may indicate interaction-based functionality switching. We further report new RBP–RBP networks along the essential processing steps of RNA metabolism. Finally, we use our screen to reveal how pathogenic mutations of splicing factors rewire their interaction networks, providing possible explanations for the observed splicing defects.

## Results

### RBPome library assembly, screen, and validation

We assembled a screen library of 978 mammalian RBPs and 76 MAPs (“RBPome library”, Table S1), and screened the library using the rec-Y2H pipeline (Figure 1A). The choice of proteins was guided by published surveys of RBPs (Hentze et al., 2018), their functional importance, and by broad coverage of the important processes in RNA biology including splicing, transport, translation, and stability control (Figure 1B). First, to ensure precision of screening, we identified auto-activating bait proteins (Figure S1A) that can give rise to false positives by screening the RBPome bait library against the empty prey vector. Detected auto-activating proteins were removed from the bait library for all following screens and detected interacting pairs with auto activators as baits were removed from final screen results. To benchmark the sampling completeness of the RBPome library screens, a subset library containing only 47 baits was assembled and screened five times against the full prey library (“H47 library”, Figure S1B, Table S1). Such a replication count saturates screens with this level of library complexity ((Yang et al., 2018), Figure S1C), and allowed us to build a reference set of interactions. The interaction score cut-off value was defined by F1-scoring (Figure S1D, (Yang et al., 2018)) against known interactions (the union of the BioGRID (Oughtred et al., 2019) and HIPPIE (Schaefer et al., 2012) databases). Comparing the true positive rate of the H47 library screen to randomized interaction matrices provided strong statistical evidence for the accuracy of the obtained screen data (Figure 1C). The RBPome screen was then benchmarked against the H47 reference set providing an F1 score-based cut-off value (Figure S1E), and showing that we reached a plateau of optimal sensitivity and specificity already after screening the RBPome library 8 to 9 times (Figure 1D). In agreement with this, we reached a plateau of newly detected interactions (Figure 1E), as well as of sampling and pair complexity after the same number of screen repetitions (Figure 1F). Overall, a large fraction of all input proteins (97.7% of baits and 98.1% of preys) were sampled in the screen (Figure S1F), which is reflected in the sampling complexity of 58.2% and a pair complexity of 77.1% (Figure 1F). This corresponds to 613,046 sampled interactions and 406,422 sampled unique interactions. As this sampling space is much larger than in previous studies (Yang et al., 2018), we devised a new interaction score calculation method (“sumIS”, see Methods) which increase sensitivity while maintaining specificity (Figure S1G). This is an important improvement of the rec-Y2H method as it now allows screening large scale libraries with this comparatively low-tech and inexpensive technique (Yang et al., 2018). Finally, we validated rec-Y2H screening results with an independent approach. We chose the NanoBRET assay (Machleidt et al., 2015) as it differs in all key points from rec-Y2H screening (interaction detection in the cytoplasm of HEK293T cells based on bioluminescence resonance energy transfer, instead of detection in the yeast nucleus based on transcriptional activation). We used modified NanoBRET bait and prey vectors (Yang et al., 2018) and found that the NanoBRET validation rate initially increased with higher sumIS thresholds, but remained consistently high above an F1 score-based cut-off of 7.1 (Figure 1G), resulting in an overall validation rate of 81.3% for interactions above cut-off which were tested by NanoBRET (Figure S1H). This also confirms that F1 scoring is a reliable tool for sumIS cut-off determination, and that even interactions in the range of sumIS 4.5–7.1 contain high-quality interaction data. We have therefore included all interactions detected above a cut-off of 4.5 in the screen results table of this screen (Table S2) but decided to work only with interactions of sumIS ≥7.1 throughout this article to ensure maximum accuracy. The strength of the interaction signals are correlated between the NanoBRET and rec-Y2H assays, providing information on confidence and reproducibility of interactions: NanoBRET MBU counts were higher among interactions tested positive in rec-Y2H (Figure S1I), and sumIS values were higher among interactions tested positive in NanoBRET (Figure S1J). Computing an enrichment score for interactions detected among and between RBPs participating in different RNA metabolic processes (Figure 1H) shows that we detect an enrichment of interactions between the same and related processes (e.g. mRNA 3’ processing and nuclear export), which support that the detected new complexes are likely of physiological relevance. Overall, our RBPome interaction screen adds 1994 new RBP interactions compared to available resources (here defined as the union of BioGRID, HIPPIE and HuRI (Luck et al., 2020) databases, Figure 1I). This and the high orthogonal validation rate set our new dataset apart from existing resources.

**Figure 1:**
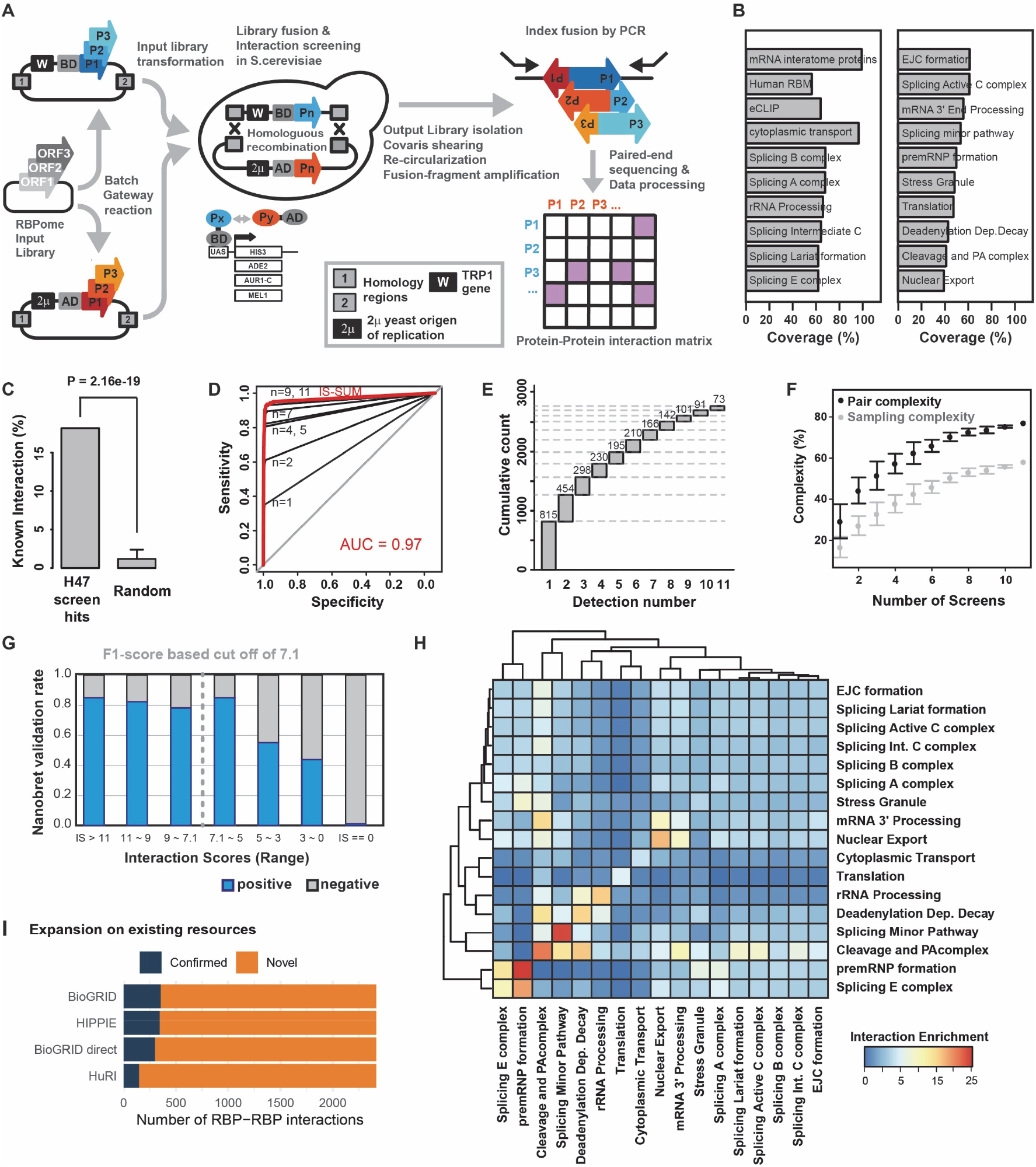
RBPome library assembly, screen and validation. **(A)** Schematic of rec-Y2H matrix screening workflow. RBPome bait and prey libraries were transferred in the respective screen vectors by batch Gateway cloning. After linearization, both vector pools are transformed together into yeast. After selection for vector fusion by homologous recombination and protein interaction selection by growth on selection media, output libraries are prepared, processed to produce NGS sequencing libraries, sequenced and interaction matrices are calculated. **(B)** Composition of the RBPome screen input library. **(C)** Benchmarking of the H47-calibaration library against the union of HIPPIE and BioGRID databases. The interactions detected above F1-score based cut-off (Figure S1D), contained a significant higher fraction of known interactions than all random resamples. **(D)** Benchmarking of the full RBPome library screens against the H47 calibration screen. After nine repetitions of the full RBPome library screen testing >1M possible interactions, sensitivity and specificity are maximized. **(E)** Cumulative number of protein-protein interactions detected with each RBPome library screen. **(F)** Pair and sampling complexity. Sampling complexity measures to which extend all possible interactions were sampled, which includes redundant pairs in both possible orientations. Pair complexity measures whether a possible interaction was sampled at least once, irrespective of the orientation. **(G)** The NanoBRET-validated rate sorted by categories. rec-Y2H scores were binned into 7 different groups. Above the cut-off (IS = 7.1), more than 80% of tested interactions were validated with NanoBRET. The validation rates were correlated with rec-Y2H interaction scores. **(H)** Enrichment matrix of interactions detected grouped by mRNA metabolic processes. **(I)** Fraction of new and confirmed interactions relative to their annotation in interaction databases and a human proteome-scale interaction screen (HuRI).

### Direct binary RBP interactions predict combinatorial RNA binding

We hypothesized that RBPs forming complexes in our rec-Y2H screen should also show an increased tendency to bind the same RNAs, or even to bind in proximity along the primary sequence (Figure 2A), if they act as complexes *in vivo*. To test this, we intersected the rec-Y2H screening PPI data with published eCLIP data (Sloan et al., 2016, Table S3) on RBP binding sites within RNAs. As positive controls, we used 5 pairs of RBPs that were detected in our screen and that had a least 5 independent literature sources in BioGRID confirming direct interaction (U2AF1-U2AF2; FMR1-FXR2; HNRNPK-QKI; RBFOX2-QKI; and SFPQ-NONO). In most cases, these pairs are also known to act as heterodimers and bind RNA as a complex (Fernandez et al., 2015; Li et al., 2018; Petti et al., 2019; Wu et al., 1999). As negative controls, we used randomly generated sets of RBP pairs of the same size as the screen hits. 94 of the RBPs we screened had published eCLIP data available, and rec-Y2H screening revealed 71 interactions between these RBPs (Table S2). This more than doubles the number of pairs previously available (Figure S2A). To quantify RNA target set similarity between RBPs, we employed the Jaccard index (Figure 2B), and to investigate whether an RBP of interests tends to bind given the presence of another RBP, we calculated the conditional probability of co-binding (Figure 2C) for all three groups of pairs (positive, identified interactions, and random pairs). To assess statistical significance, we calculated a resampling p-value using 10,000 iterations of random pair sampling, with the newly identified interactions always scoring higher than random pairs for both (i.e. p<0.0001). The positive controls scored highest in both cases. This analysis clearly shows that on average, RBP pairs detected by rec-Y2H screening share more RNA targets and have a higher probability of binding given the binding of their interaction partner than random pairs, thus providing initial support to the hypothesis that RBPs we found to interact indeed bind RNAs as complexes.

**Figure 2:**
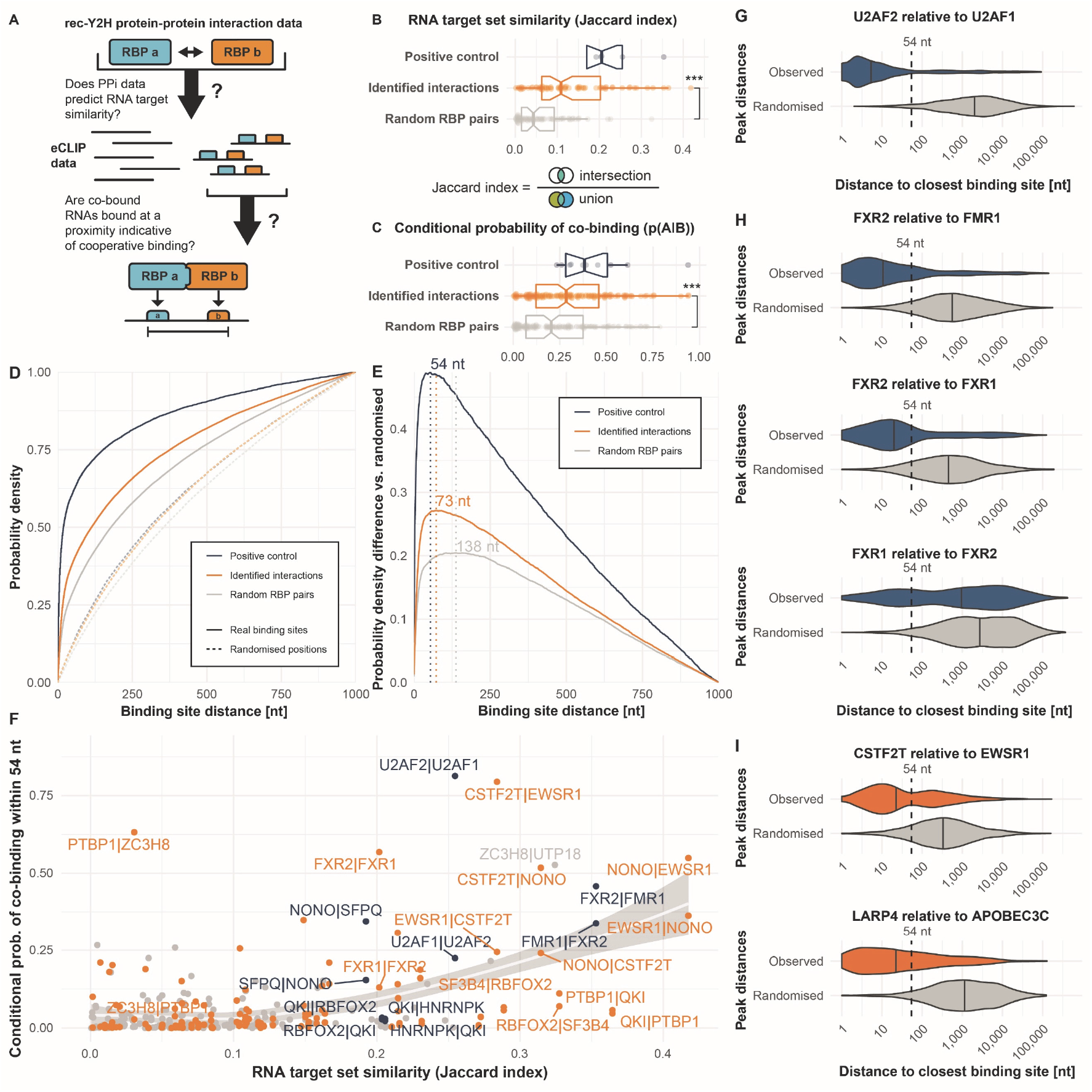
Protein–RNA interaction data provides support for complexes of RNA-binding proteins. **(A)** Binding sites of non-interacting pairs of RNA-binding proteins (RBPs) may be distant from one another, while binding sites of interacting RBPs should be in proximity. **(B)** Interacting RBPs display greater target set similarity than random RBP pairs, as measured using the Jaccard index of their RNA target sets. **(C)** Interacting RBPs display a greater conditional probability of co-binding (RBP A binding, given RBP B binding) than random RBP pairs. **(D)** Binding site distances tend to be smaller for interacting RBPs than for random RBP pairs, and they tend to be far smaller than between randomly positioned binding sites within the same RNAs (dashed lines). **(E)** The difference between the observed distances and the simulated distances between randomized sites within the same RNAs is maximized at ≤54 nt for positive control cases. This suggests confident separation between cases of binding as a complex or as individual RBPs using this threshold. **(F)** Conditional probability of RBP–RBP co-binding within ≤54 nt, plotted against the RNA target set similarity of the RBP pair. Each pair is plotted in both conditional probability orientations (p(A|B), i.e. A given B, and p(B|A), i.e. B given A). **(G)** Probability density “violin” plots showing observed binding sites compared to the expectation from randomized positions (grey). Medians are indicated by vertical stripes. When investigating U2AF1 binding sites, U2AF2 is generally found binding very closely to it (≤54 nt), demonstrating strong dependency of U2AF1 on U2AF2. **(H)** Similarly, when investigating FMR1 and FXR1 binding sites, FXR2 is generally found binding very closely to these (≤54 nt), indicating their dependency on it. However, the inverse is not true: when investigating FXR2 binding sites, the nearest FXR1 binding site can be distant, indicating independent binding by FXR2 (potentially with its alternative partner, FMR1). (**I**) Among RBP–RBP interactions newly identified in our screen (orange), EWSR1 binding sites tend to be close to CSTF2T binding sites, indicating some dependency on CSTF2T. Likewise, APOBEC3C binding sites tend to be close to LARP4 binding sites (≤54 nt), indicating a degree of dependency on LARP4.

### Newly discovered RBP pairs bind RNAs in immediate proximity

To test which RBP pairs bind in close enough proximity to allow adjacent RNA binding as complex, we next analyzed the distance of eCLIP-derived RNA binding sites for the same three groups of RBP pairs (positive controls, identified interactions, and random pairs) and generated cumulative density functions (CDFs) for these (Figure 2D). As matched controls, we included randomized binding positions within the same set of RNAs for each group, thereby maintaining the binding density, number, and length distribution of RNAs (see Methods). Since proximity along the primary sequence is only likely to be informative for potential binding as a complex within a reasonable distance, we discarded binding site combinations more distant than 1000 nucleotides. The positive controls and the screen hits had a considerably higher fraction of binding events at closer distances compared to random RBP pairs, with their CDFs rising more quickly, while the matched controls with randomized binding positions were close to a fully random distribution at the diagonal. To derive a characteristic binding site distance threshold below which RBPs are most likely to bind RNA as complexes, we determined the point of maximal difference between the observed binding site data and the matched randomized controls. This was done by subtracting the randomized density function from the observed one (Figure 2E). For the positive controls, we found that a binding site distance threshold of 54 nt produces the highest excess of observed co-binding events compared to those expected by chance. When calculating this metric for the identified interactions data set, the optimal threshold was only slightly higher at 73 nt. For the random pairs we calculate a clearly higher maximal difference to randomized binding data at 138 nt. Taken together, our analysis demonstrates that RBP pairs detected with rec-Y2H screening have an increased probability to co-bind RNA adjacently, which is indicative of binding as a complex. Interestingly, several of the random RBP pairs initially introduced as negative controls scored highly in terms of their target set similarity (Jaccard index), co-binding probability, and binding site distances (Figure 2B&C). These are likely false-negatives that rec-Y2H screening could not detect or detected interactions that resampling generated by chance. To test this, we analyzed 40 random pairs using NanoBRET. Indeed, we detected 12 positive interactions (Table S4), suggesting that the difference to true negative pairs would be higher than we calculated here.

To illustrate the most striking examples of detected RBP complexes, we plotted their conditional probability (“A binding given B binding”) of co-binding within 54 nt against their target set similarity (Figure 2F). Pairs falling into the upper right portion of the plot have highly similar RNA target sets, as well as a strong dependency of the left-hand protein to be present where the right-hand one binds, i.e. obligate co-binding. As an example, CSTF2T has a remarkably high probability of binding adjacent to EWSR1 (CSTF2T|EWSR1). This indicates a strong dependency of EWSR1 on CSTF2T to be present. The inverse is less true: CSTF2T appears less dependent on EWSR1’s presence (see EWSR1|CSTF2T). Meanwhile, pairs in the upper left portion display lower RNA target set similarity while still displaying high conditional probabilities of co-binding, i.e. while the right-hand protein is highly dependent on the other, the left-hand protein appears to bind additional targets, either independently or as part of other complexes. Figures 2G–I illustrate the distribution of minimal binding site distances of four identified interactions classified as known positive controls (Figure 2G&H) and two newly detected pairs (Figure 2I). These cases show a remarkably close median binding site distance ranging from 4–20 nt, compared to their matched randomized binding site controls, which show medians between 338–2830 nt. An exception is the FXR1-FXR2 pair which also shows a low conditional co-binding probability (Figure 2F). Of note, these asymmetries in conditional co-binding probability could also at least partially result from differences in eCLIP data quality.

In summary, this analysis shows that compared to randomly generated RBP pairs, pairs detected by rec-Y2H screening bind significantly more similar sets of RNAs, and have a higher probability to bind their targets in proximity at or below 73 nt, suggesting that they act as complex.

### The choice of RBP complex partner directs the site of RNA interaction

The biological functions of RBPs can be linked to the RNA regions they bind, such as intron boundaries or the 3’ UTR region (Mukherjee et al., 2019b). To which degree RBPs can adapt their function depending on their combination in different complexes, is an interesting and intensely debated question (Achsel and Bagni, 2016; Quattrone and Dassi, 2019; Sternburg and Karginov, 2020)(Figure S3A). To show an overview of RBPs with strong evidence of binding adjacent to each other, we first created a network of RBP pairs (Figure 3A) that bind significantly more frequently in proximity (≤54 nt) relative to randomized binding positions according to a combination of two resampling-based statistical tests. Out of the 71 screen hit RBP pairs for which eCLIP data was available, 53 showed significant proximity binding according to at least one test (Figure S3B and Table S5), and 30 showed significant proximity binding according to both tests (Figure 3A). To investigate whether the pre-mRNA region preference of proximity-binding RBP pairs deviates from the preferences of the individual RBPs, we computed meta-pre-mRNA profiles for mRNAs bound by the individual RBPs (Figure 3B, left panel) and by a subset of pre-mRNAs which are bound by RBP pairs in proximity (Figure 3B, right panel). Clustering of both heatmaps showed two distinct behaviors of RBPs. On one hand, there are conservative, presumably uni-functional RBPs which exhibit the same pre-mRNA binding profile alone and on pre-mRNAs which they co-bind in proximity with an interaction partner (Figure S3C). Examples are nuclear heterodimers such as U2AF1 and U2AF2 (Figure 3B, C&D) or the NONO-EWSR1-SFPQ-CSTF2T complex (Figure 3A, B & Figure S3D) which are intron binders alone and on co-bound transcripts. On the other hand, proteins such as PTBP1 alter their binding profiles depending on the interaction partner. Alone or on transcripts co-bound with the zinc-finger protein ZC3H8, PTBP1 is a clear intron binder (Figure 3B, E & Figure S3E), while on transcripts co-bound with PCBP1, intron and 3’UTR binding is equally strong (Figure 3E) and its individual binding profile correlates poorly with the one found on co-bound transcripts (Figure 3F). While the homologues PCBP1 and 2 have identical binding profiles on pre-mRNAs co-bound with IGFBP2 (Figure S3F), PCBP1 exhibits diverse profiles depending on its many interaction partners (Figure 3G) and also its interaction partners alter their profiles on RNAs co-bound with PCBP1. HNRNPK, for instance, shows a strong preference for intron-binding, but on RNAs co-bound with PCBP1 it shows a similar high density in 3’UTRs (Figure 3B & G). Both protein bind C-rich sequences. However, while HNRNPK is mostly associated with pre-mRNA processing and PCBP1/2 is also involved in translation and mRNA stability regulation, the observed pre-mRNA binding region preference shift of HNRNPK could indicate that it also acts as complex with PCBP1 in 3’UTRs to control later processes in the life of an mRNA. Because PCBP1 shuttles between the nucleus and the cytoplasm, it is perceivable that it encounters different interactions partners in different compartments which redirect its activities. Indeed, there is a higher transcript binding profile similarity between PCBP1 and its nuclear interaction partners (HNRNPK and PTBP1) than among profiles with shuttling or cytoplasmic RBPs. Overall, our analysis shows that there is a widespread potential for RBPs to adjust their functions depending on their complex partners (Figure 3H); when comparing Pearson correlation values of pre-mRNA binding profiles for all RBPs with more than one interaction (15), 9 RBPs form at least one complex which shows an only moderate positive relationship (p≤ 0.7) to their individual binding profile. We expect that the analysis we develop here will reveal the full breadth of this phenomenon as more eCLIP data will become available in the future.

**Figure 3:**
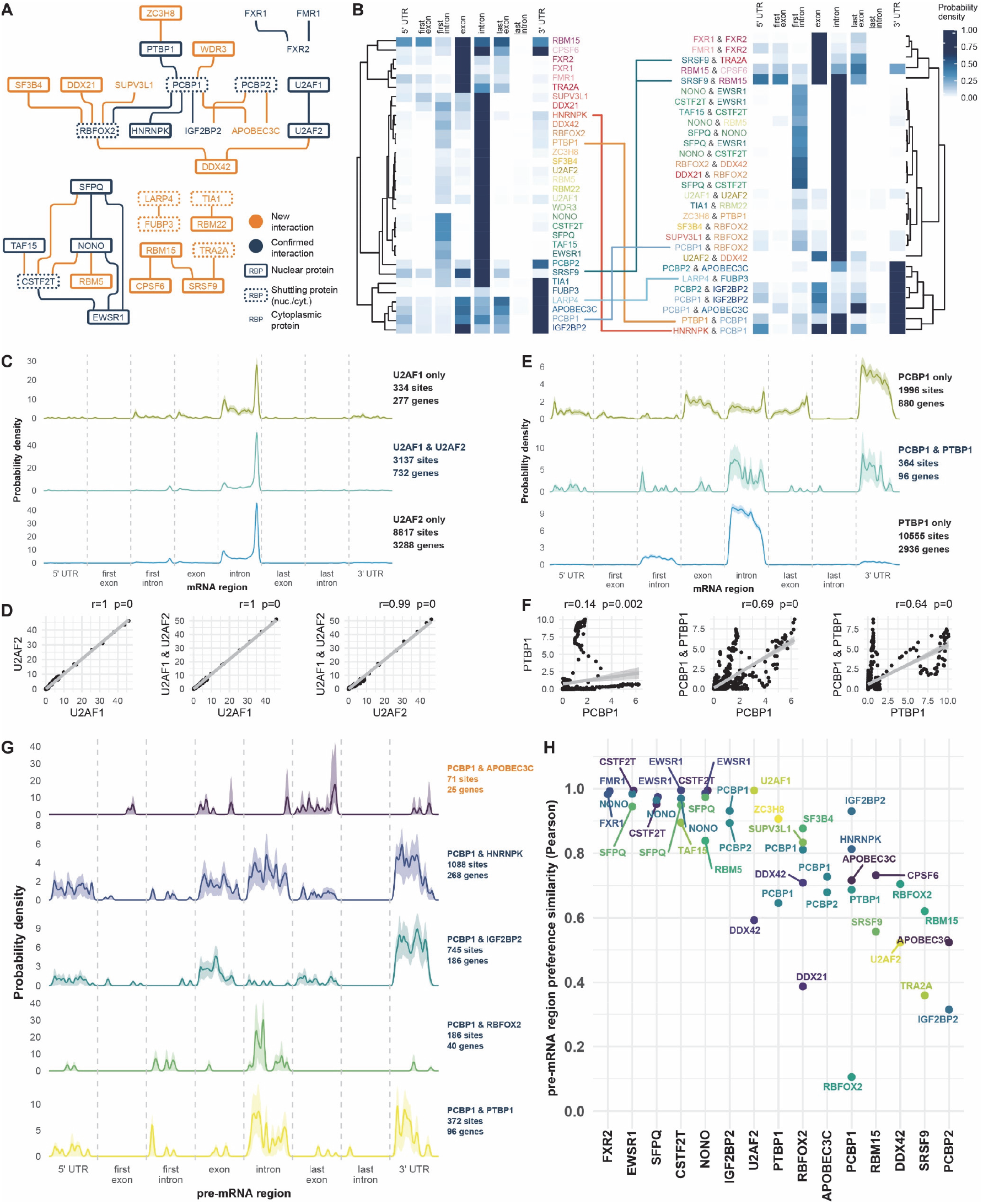
Complexes can be guided to different mRNA regions by their members. **(A)** Network graph of RNA-binding proteins with eCLIP protein–RNA interaction data. Only pairs with strong statistical support (see Methods) for significant binding in proximity within 54 nts in at least one pair orientation are shown. Subcellular localization is indicated by gene symbols framed either by a continuous border (nuclear proteins), a dotted border (shuttling, i.e. sometimes nuclear), and no border (always cytoplasmic). Interactions newly identified in our screen, and the proteins newly joined by them to the network, are highlighted in orange. **(B)** Hierarchically clustered heat maps showing the mRNA region preferences of the individual RBPs (left) and the RBP pairs from the panel A network (right), based on published eCLIP data. For pairs, only binding sites in proximity close enough to be indicative of binding as a complex (within 54 NTs) were included. The color coding is maintained for each RBP, and manually added lines tracing individual RBPs to certain pairs indicate cases where the pair shows a binding region pattern that is considerably different from the individual protein. **(C)** Probability density estimate plots showing the underlying data for the panel B heat map for U2AF1 & U2AF2. Genomic coordinates were mapped to pre-mRNA regions by using the architecture of the most highly expressed transcript for each gene, averaged across the two eCLIP cell lines. U2AF1 only binds independently of U2AF2 at a small fraction of sites (342 sites), and the mRNA region preferences of the complex mirror those of U2AF2 (binding in introns, and particularly at their 3’ end). **(D)** Pairwise binding profile correlation plots (Pearson’s r) of the examples shown in panel C. Each axis shows the probability density estimate for either an individual RBP or that of the pair in proximity. **(E)** Probability density plots of eCLIP profiles of pre-mRNAs bound by PCBP1 or PTBP1 alone and in proximity. **(F)** Pairwise binding profile correlation plots of the examples shown in panel E. **(G)** Probability density plots of different subsets of pre-mRNAs bound by PCBP1 in proximity with alternative co-binding partners (putative complexes). **(H)** Pearson correlation coefficients of pre-mRNA binding profiles. Only RBPs with more than one detected interaction partner are shown. The colored dots show the Pearson correlation values of the pre-mRNA binding profiles of RBP pairs vs the profile of the individual RBPs shown along the x-axis.

### RBP complexes recognize specific di-motifs

For many RBPs, multiple possible sequence motifs have been reported, and these can sometimes differ significantly (Giudice et al., 2016). We speculated that RBPs might use different motifs depending on the complex partner they bind RNA with. In this way, combinations of different motifs at the correct spacing could guide the specific binding of RBP complexes. To test this idea, we asked two questions (Figure 4A): 1) Do we detect different motifs within eCLIP peaks where RBPs bind without a specific interaction partner than in regions where the two RBPs bind adjacently? 2) Do RBPs prefer different motifs depending on the interaction partner with which they bind adjacently (within 54 nt)?

**Figure 4.**
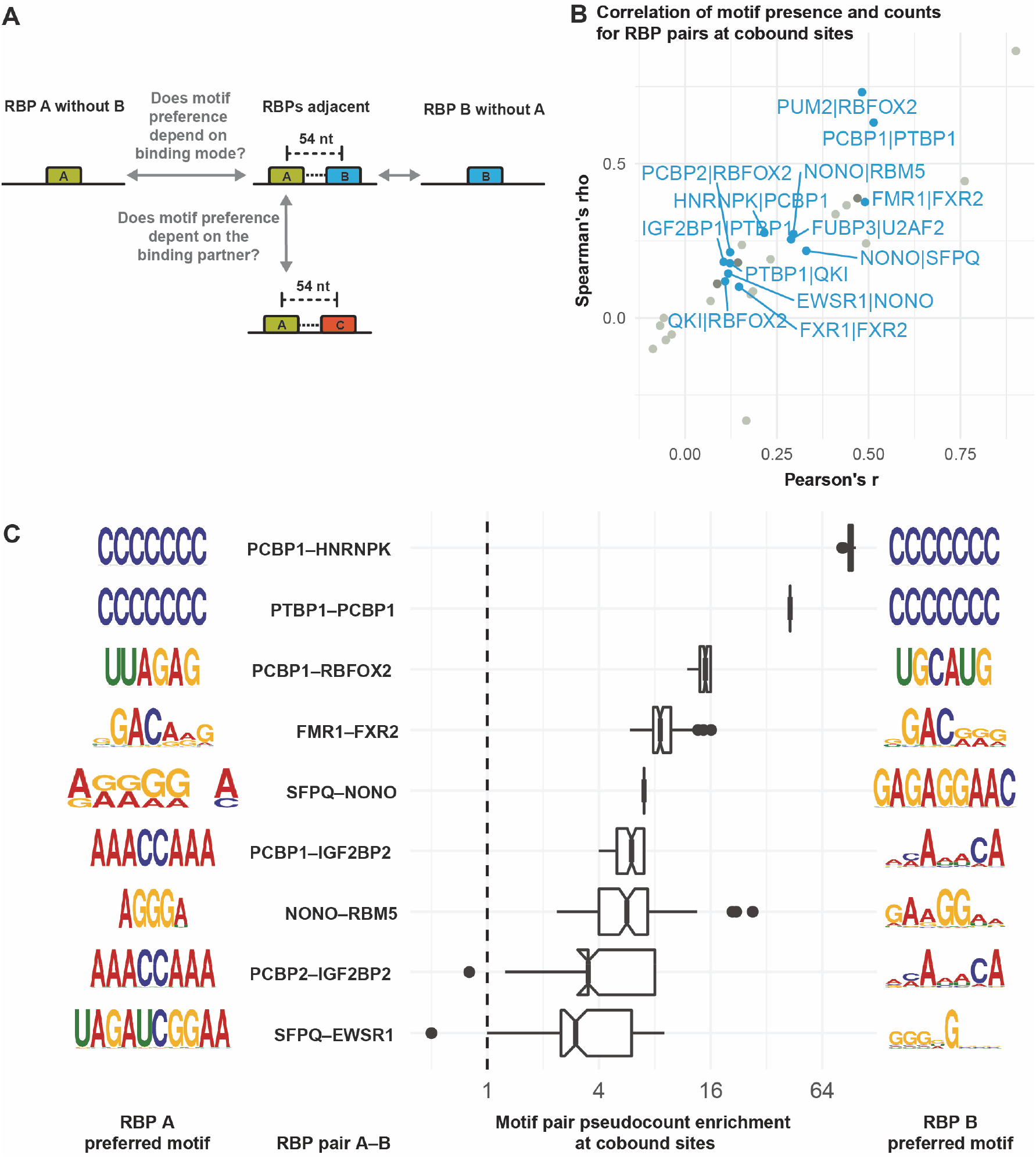
RBP-pair di-motif preference. **(A)** Motivation for this analysis. **(B)** Correlation of motif presence/absence (Fisher’s exact test) and motif counts (Spearman and Pearson) at co-bound sites, i.e. where interacting RBPs bind in proximity (≤54 nt). RBP pairs for which all tests are significant at p<0.05 are labelled. The Fisher test indicates whether having a motif at all is correlated between the two RBPs, and it takes precedence for the color coding. Blue dots: all tests significant, dark grey dot: insignificant motif count correlation, light grey: insignificant motif presence correlation. **(C)** For each RBP pair, the preferred combination of motifs for RBP A (left) and RBP B (right) when both bind in proximity is shown as a sequence logo. The box plots show the pseudocount ratio indicating how frequently a given motif combination occurs when two RBPs bind in close proximity as compared to an independent-binding background derived from target genes where only one of the RBPs binds.

For each RBP pair with known motifs, we first assessed whether the presence of any known motif for one RBP within its binding region correlates with motif presence for the adjacent RBP in its own binding region using Fisher’s exact test. Since we observed that many binding regions contain multiple copies of a given motif, we also tested whether the overall numbers of motifs within binding regions are correlated for a given RBP pair. We find that for 13 out of 33 interacting RBP pairs with known motifs that bind in close proximity, motif presence as well as motif count are significantly correlated at adjacently bound sites (Figure 4B).

Next, we investigated motif pair combinations, i.e. whether specific known motifs tend to occur together when two RBPs bind adjacently. We compared these motif pair frequencies to a background in eCLIP peak regions of genes where only one of the two interacting RBPs binds (Figure 4C). For 9 RBP pairs with significant evidence of co-binding in proximity (Figure 3A), we find that certain motif combinations are at least 3-fold enriched at co-bound sites. For each RBP, the most enriched motif is shown, with a single representative being chosen in case of ratio ties (see Figure S4A for an extended version showing all top-enriched motifs including ties). When comparing the motif of choice of the same RBP when it is in complex with different interaction partners (Figure 4C), we find that the preferred motifs deviate strikingly (e.g. NONO with EWSR1 vs. NONO with SFPQ, or PCBP1 with either of HNRNPK, RBFOX2, or IGF2BP2). This analysis shows that RBPs use different motifs when they bind adjacently compared to binding in isolation, and also that the sequence motif preference of at least some RBPs may depend on their specific interaction partner. We expect that the depth of this analysis will be extended significantly as more eCLIP data becomes available and more motifs are discovered.

### Rec-Y2H screening reveals new RBP-networks along the life of an mRNA

To next illustrate how rec-Y2H screening contributes to the understanding of different processes of mRNA metabolism, we created interaction networks filtered by RBPs belonging to different process (Table S6), based on the reactome database (Jassal et al., 2020). Processes not listed in reactome were based on other databases (Youn et al., 2018) or a literature curated list of involved proteins as in the case for cytoplasmic mRNA transport. A limitation of Y2H-based screens is that all screened proteins are fused to nuclear localization sequences, which can produce false-positive interactions between proteins that naturally would not be present in the same compartment. Therefore, we first tested which fraction of interactions were detected between proteins of the same compartment (nucleus, cytoplasm), and found that screen hits have significant higher interactions between RBPs of the same compartment compared to randomly generated pairs (61.6% vs 21.6%) (Figure S5A). Of note, as only primary locations annotated in the Human Protein Atlas (Thul et al., 2017) were used, even some interactions between proteins classified differently could be meaningful due to e.g. shuttling activity. Our analysis revealed intriguing new interactions along many important steps of an RNA life. For all classes of RBPs we confirm known interaction and we discover new interactions, with the highest fraction of new interactions identified for processes taking place in the cytoplasm such as cytoplasmic mRNA transport and stress granule formation (Figure 5A). For other nuclear processes such as splicing (Figure 5B & Figure S5B&D), with the exception of the minor splicing pathway (Figure S5C), we have a higher ratio of confirmed interactions, potentially because these processes are well investigated already. The newly detected interactions between the major EJC component MAGOH, the 3’end processing hydrolase NUDT21 and the components of the cleavage factor Im complex (CPSF7, Figure 5C), could hint towards a new link for the coordination of nuclear export, translation regulation and mRNA processing. Concerning nuclear export (Figure 5D), we detected new, direct interactions between the principal mRNA export factor NXF1 and several SR proteins. These interactions are considered transient and stabilized by RNA as indicated by their RNAse sensitivity (Müller-McNicoll et al., 2016). The fact that we do detect these interactions is a good indication that our methods can detect weaker, dynamic interactions that are lost by RNAse treatment during AP-based techniques. A major open question of mRNA metabolism is how mRNPs are coupled to cytoplasmic, microtubule-based motor proteins. Apart from a few reconstituted complexes (Baumann et al., 2020; Heym et al., 2013; McClintock et al., 2018; Sladewski et al., 2013), data on RBP-motor coupling are mostly based on pull-down experiments (Charalambous et al., 2013; Dictenberg et al., 2008; Kanai et al., 2004; Mallardo et al., 2003) which can only inform about physical but not direct interactions. Here we show that two factors that have been previously indirectly connected to cytoplasmic mRNA transport (NXF1 (Pocock et al., 2016), the TSN-TSNAX complex (Chennathukuzhi et al., 2003; Severt et al., 1999)) indeed directly interact with kinesins and a myosin (TSNAX), or in the case of NXF1 even with dynein cargo adaptors (BICD2) and microtubule plus end tracking proteins as EB3 (Figure 5E). The detected new interactions of NXF1 are at first surprising as NXF1 is thought to rapidly re-enter the nucleus after mRNA export (Müller-McNicoll and Neugebauer, 2013) and could hence not have a function in cytoplasmic mRNA transport; however NXF1 has recently been detected in cytoplasmic polysomal fractions (Botti et al., 2017) and in the neurite-fraction of induced neurons (Zappulo et al., 2017) which opens the possibility that NXF1 guides the journey of some mRNAs further, through the cytoplasm. Lastly, while the composition, biophysical and biocheMICal processes of stress granules and processing body formation is intensely studied, the direct molecular interactions between their constituents are poorly understood. Here we provide a first direct interaction network of proteins found in stress granules, processing bodies or both (Figure 5F), identifying 92 new interactions among these proteins. We have used only tier-1 proteins (Youn et al., 2018) and show only interactions above an sumIS threshold of 10, for reason of presentability. All detected interactions among stress granule (tier-1) and processing body proteins are listed in the Supplemental Table S6. Interestingly, newly identified interactions connect the m6A-reading proteins in YTHDF1 and 3 to core components of stress granules as QKI and RBFOX2. This is of high interest as recently it was found that m6A-modified mRNAs are enriched in stress granules and that YTHDF1 and 3 are essential for their formation (Fu and Zhuang, 2019). Regarding factors involved in rRNA processing (Figure S5E), we detect new interactions between exosome components (EXOSC7&8) and ribosomal subunits (RPL0,3,21 and RPS28) which indicate that direct interactions between ribosomal proteins and these exosome components could play a role in rRNA processing. Finally, all but one detected interaction involved in regulation of mRNA stability (exosome and LSM complexes) were previously known (Figure S5F), which illustrates that rec-Y2H screening can precisely capture the architecture of protein complexes; the only new interaction detected between PABPC1 and EXOSC8 could point towards a function of PABPC1 in recruiting the exosome complex to the mRNA 3’end. A Cytoscape (Saito et al., 2012) file with all interactions detected above the sumIS thresholds of 4.5 or 7.1 and NanoBRET validations is provided (Data S1), which allows to explore more subnetworks of interest.

**Figure 5.**
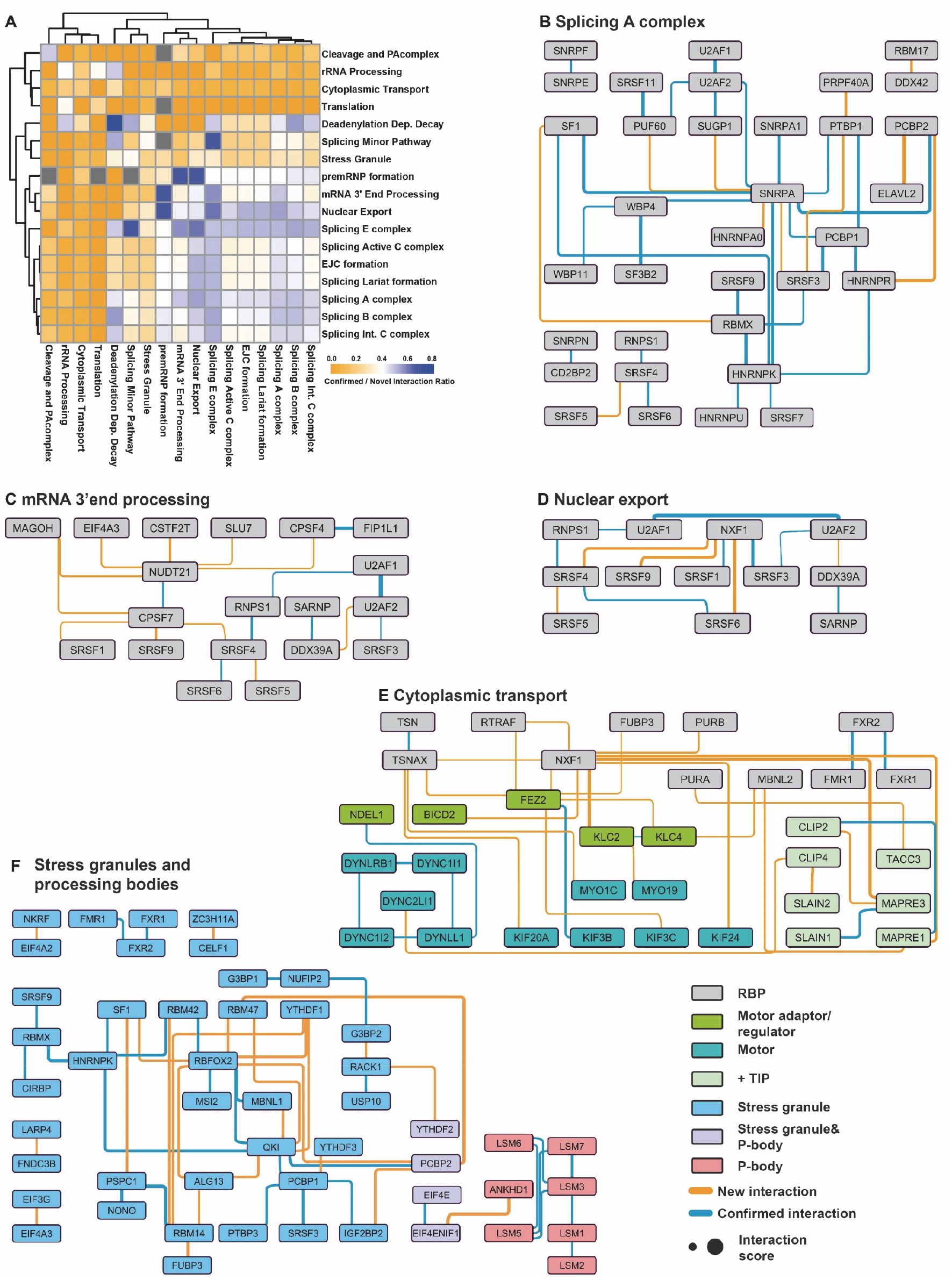
New interactions along the life of an mRNA. **(A)** The matrix illustrates the ratio of confirmed and new interactions detected among proteins belonging to the respective mRNA metabolic processes. **(B-G)** The process classification is based on the reactome database with the exception of stress granules which were defined based on the RNA granule database and cytoplasmic transport, which is based on literature-curated information. For the selection of the proteins involved in cytoplasmic mRNA transport (F), all motor proteins and microtubule binding proteins were included along with all RBPs which were related to cytoplasmic mRNA transport in the literature. For the definition of stress granule and processing body constituents all proteins classified as tier-1 at the RNA granule database were chosen. Due to the amount of detected interactions, only interactions above a sumIS of 10 are shown in (G). A full list of interactions in stress granules can be found in Supplemental Table S6.

### Cancer associated RBM10 mutations impair interactions with splicing factors

Taking advantage of our RBPome-scale screening library we next decided to test if and how disease relevant mutations in splicing factors affect the topology of splicing networks. To increase confidence in mutant screening results, which often resulted in a reduction or increase of interactions scores, we had included all mutants in both the full RBPome library and the H47 library screen. Comparing both screen matrices shows that the results of both screens (Table S7) are highly similar (Figure S6A & S6B) and that even the direction in which interaction scores change in response to mutations are significantly correlated (Figure S6C). We included mutants of the tumor suppressor RBM10 in our screen library affecting different regions of the protein (Figure 6A). The V354E and Y580F mutants were originally detected in lung cancer patients (Bechara et al., 2013; Imielinski et al., 2012) and the V354E mutation was shown to abolish NUMB alternative splicing activity of RBM10 (Bechara et al., 2013). Correct NUMB splicing leads to exon-skipping and the production of a NUMB variant which prevents notch-receptor accumulation, a critical event promoting lung-cancer progression (Misquitta-Ali et al., 2011). Interestingly, the V354E mutations is not affecting the RNA binding ability of RBM10 (Hernández et al., 2016). This is surprising as the mutation lies in the second RRM domain of RBM10 (Figure 6A). We hence hypothesized that this mutation must impair the ability of RBM10 to form functional complexes with other components of the splicing machinery which then causes the observed perturbation in alternative splicing. This is not too unlikely as RRM domains were shown before to mediate protein-protein interactions (Maris et al., 2005). To test this hypothesis, we included two more RRM2 mutations (I316F & R343G), predicted by FoldX (Guerois et al., 2002) to disrupt the folding of the RRM2 domain, into our screen library. This is expected to have similar effects on RRM2 mediated protein interactions (Figure 6B). Our analysis shows that the V354E mutation leads to a striking loss of interactions with three key alternative splicing factors (DDX17, PTBP1 & 2), the NEXT component RBM7, the U1-snRNP component SNRNPA and two RBM12 variants (Figure 6C-E). An effect of PTBP2 on alternative NUMB mRNA splicing is known (Licatalosi et al., 2012), which possibly hints towards a joined function of RBM10 and PTB2 in this process. The two control mutations that disrupt RRM2 folding had remarkably similar effects on RBM10 interactions (Figure 6C & E), supporting the idea that the RRM2 domain of RBM10 coordinates health relevant splicing events through its protein-protein interactions with other splicing factors.

**Figure 6.**
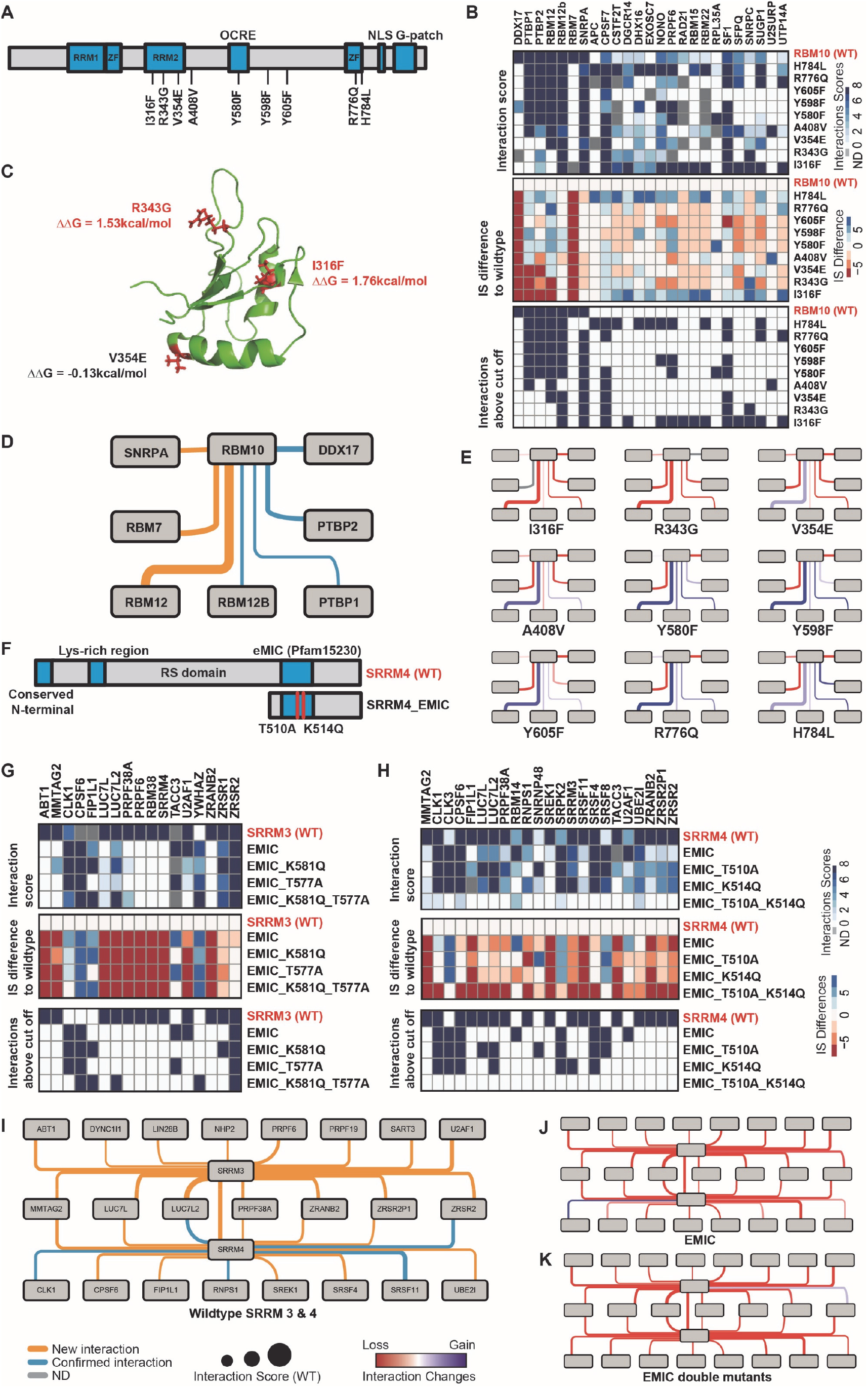
RBM10 RRM-domain mutations disrupt splicing networks. **(A)** Schematic representation of the RBM10 domain architecture and mutants tested. **(B)** Interaction matrix of wildtype RBM10 and mutants. Upper panel: Interaction score (sumIS) of all tested combination. ND (grey) represents the non-detected cases which can either indicate a complete loss of interaction (true-negative) or false-negatives caused by incomplete library sampling (false-negative). Middle panel: Difference of interaction score between wildtype and mutants. A reduced interaction score is shown in red, an increased interaction score in blue. Lower panel: cut-off filtered representation of the matrix shown in the upper panel. Only interactions scoring higher as the F1-score determined cut-off of 7.1 are shown. **(C)** Crystal structure of the RBM10 RRM2 domain. The three mutations affecting this domain are highlighted in red and the FOLDX-computed delta free energy for domain destabilization is shown for each mutant. A positive change of Gibbs free energy indicates mutations that are more likely to destabilize the structure. **(D)** Interaction network of wildtype RBM10 showing only interactions above cut-off. **(E)** Interaction networks of RBM10 mutants with the same topology as the wildtype network shown in (D). Lost interactions are shown in red and gained interactions are shown in blue. Not detected interactions are shown in grey. **(F)** Schematics of the domain architecture of SRRM4 and mutants tested in our screen. **(G)** Interactions matrices showing detected direct interactors of SRRM3 and mutants. Upper, middle and lower panels as in (B). **(H)** Interactions matrices showing detected direct interactors of SRRM4 and mutants. Upper, middle and lower panels as in (B). **(I)** Interaction network showing the common and individual interactions of the paralogs SRRM3 and 4. **(J & K)** The network topology is the same as in (I). The color of the connecting lines indicates gain or loss of interaction score relative to the wildtypes.

### SRRM3 and SRRM4 have similar but not identical interaction networks

As a second example, we investigated the interaction network of the SRRM3 and 4 splicing factors. Both proteins are orthologues with 30% sequence identity (Torres-Méndez et al., 2019) and have partially redundant function in alternative splicing of micro exons. While the interactome of SRRM4 has been well studied by AP-MS (Gonatopoulos-Pournatzis et al., 2018), it is not understood to which extend SRRM3 and SRRM4 share interactors. Also, to date there is no evidence for direct interactors of the two proteins and to which extend they depend on the eMIC domain (Figure 6F), which has been identified as a crucial hub, mediating protein interactions with early spliceosome factors (Torres-Méndez et al., 2019). Our data reveal a direct interaction between SRRM3 and SRRM4, suggesting a potential for heterodimerization. This interaction is not detected for the eMIC-fragment of SRRM3 (Figure 6G-K), indicating that heterodimerization does involve the eMIC domain of SRRM4 but not of SRRM3. SRSF11 and RNPS1 were identified as important interactors of SRRM4 via AP-MS. All three factors are crucial for neuronal micro exon-splicing and co-incubation of these factors with SRRM4 increases their affinity to RNA (Gonatopoulos-Pournatzis et al., 2018), suggesting cooperative binding. Our screen now shows that these factors can indeed directly interact and that these interactions do not depend on the SRRM4 eMIC domain. While seven out of 15 interactions detected both for SRRM3 and 4 are shared (Figure 6I), most of these interactions were not found to depend on the c-terminal eMIC domain (Figure 6I-K). Of note, in the case of SRRM4, introduction of two loss of function mutants (Torres-Méndez et al., 2019) inside the eMIC domain caused a complete loss of the SRRM4 interactions and all other interactions including SRRM3, while this effect was less pronounced in the case of SRRM3 (Figure 6G & H), which indicates a functional diversification of the eMIC domains in SRRM orthologues.

## Discussion

RBPs are a large class of proteins that regulate the basic aspects of RNA metabolism. However, their potential to form complexes with distinct functions is not explored systematically. In this study, we focused on the discovery of direct interactions among mammalian RBPs and found a wealth of novel high-confident interactions, which are a useful resource to understand how RBPs act together. We further improved the analysis pipeline of the rec-Y2H screen to enable screening of large-scale libraries. This is a significant advance, as relative to other large-scale matrix screening methods, rec-Y2H is comparatively resource efficient (Yang et al., 2018).

The RNA binding motifs of many RBPs are short and common, suggesting that combinatorial RNA recognition by RBP complexes would be an elegant mechanism to increase affinity, specificity and the regulational potential. RBPs interacting in rec-Y2H screening do not only show an increased tendency to bind the same RNAs, they also bind these RNAs in much closer distance compared to both, randomly generated RBP pairs or randomized binding positions. This indicates that interacting RBPs bind the same RNAs as complex, potentially carrying out specific functions of RNA metabolism together. We do detect that a major fraction of RBPs can alter their mRNA binding region preferences depending on their interaction partner (Figure 3H), perhaps indicating that one of the partners directs the complex towards certain targets. It is possible that complexes could enable binding of different, potentially lower-affinity motifs. This is highly intriguing as it indicates that a significant fraction of multi-functional RBPs exist that show context-dependent activities. RBP co-binding is likely guided by specific motifs, placed at the right distance. Indeed, we find that sometimes specific motifs are preferred when RBPs bind adjacent to a specific interaction partner, and that these appear to differ for different partners (Figure 4C). We propose that the motif choice by two interacting RBPs composes novel di-motifs that might have a widespread function in guiding specific RBP-RNA interactions — an idea we will be able to test as more eCLIP data becomes available.

eCLIP data is an average over a large number of cells and RNAs. It is hence not possible to determine whether the same RNA molecule is bound by a given RBP pair at the same time. Our PPI data, however, clearly shows that the considered RBPs form complexes and the precision of our PPI data is backed up through validations with an independent method. Also, it could be considered possible that a fraction of proximity-binding events are not co-binding events but rather indication of competition. However, if this was the case, it would be difficult to envision a general scenario where competing RBPs directly interact. Of note, we regard only one possible scenario for combinatorial RNA recognition — direct RBP interaction and RNA binding as a complex at proximal binding sites. Certainly, interacting RBPs do not necessarily need to bind in proximity along the primary sequence; RNA looping could allow all kind of distances between binding sites of two interacting RBPs. However, primary sequence proximity directly implies physical proximity. Furthermore, cases exist, also for DNA-binding transcription factors, in which protein-protein interactions are weak in solution but binding to their substrate increases these interactions (Hennig et al., 2014; Jolma et al., 2015). We are likely not detecting most of these cases, even though the overexpression of proteins in our Y2H assay can lead to the detection of weaker interactions as exemplified by the weak interactions between NXF1 and SR proteins we detect. Furthermore, we showed that rec-Y2H sensitively identify protein interaction-network changes upon a single amino acid change. Hence this method can help to identify the functional consequences of protein mutations in complex diseases in the future.

Currently, eCLIP data exist for only 150 of the around 1400 known mRNA-interactome proteins (Hentze et al., 2018), while our RBPome screen covers about 80% of the known mammalian mRNA interactome with at least twice confirmed interactions. Hence, as more eCLIP data will become available in the future, the analysis pipeline developed here can be used to significantly extend the breadth of this study to generate related PPI – eCLIP datasets predicting combinatorial RNA binding at RBPome and transcriptome scale. We expect that this will significantly help to understand principles underlying protein-RNA interactions, RNA metabolism and gene expression.

## Supporting information

Supplemental Data 1 - Cytoscape file with all detected interactions

## Acknowledgments

We thank Lucas Carey, Julian Koenig, Manuel Irimia, and Sophie Bonnal for careful reading and input on the article. All sequencing was done in the CRG Genomics Core Facility.

## Author contributions

SPM conceived the project. MGC, MGS, SP performed experiments. BL, JSY, SPM analyzed the data. SPM, BL, JSY, GT wrote the article. SPM supervised and coordinated the project.

## Data and materials availability

All data needed to evaluate the conclusions in the paper are present in the paper and/or the Supplementary Materials. Additional data related to this paper may be requested from the authors. All sequencing raw-data is deposited at ArrayExpress (accession: E-MTAB-9612).

## Materials and Methods

### Screen library assembly and rec-Y2H screening

Rec-Y2H screens were performed as described in Yang et al. 2018. A detailed protocol can be found in the Supplementary section of the Yang et al. article. Minor changes were applied due to the larger size of the library screened in the present article. The number of pENTR-ORF sub-pools to build the library was adapted to cover the whole library, with a maximum of 96 ORF clones in each sub-pool. The bait library was smaller, as already known auto-activators were not included. After auto-activator pre-screening newly detected auto-activators (Figure S1A), were as well removed from the bait library, resulting in a final RBPome screen library of 1001 baits and 1054 preys. For the H47 subset screen library, 47 baits and all 1054 preys were selected. The number of yeast transformations was expanded to 48 transformation per screen, to cover as much as possible the current library, containing initially1054 × 1054 bait-prey combinations. Finally, 12 Zymoprep™ Yeast Plasmid Miniprep II columns were used to extract the yeast plasmid DNA, 20 independent R1/R2 PCR reactions and 12 P5/P7 PCR reactions for each selection media. Briefly, to assemble H1001 library, pENTR-ORF clones were either picked from the Human Entry ORFeome v8.1 collection (transOMIC) or cloned from a Human brain cDNA library. pENTR-ORFs were grown in 96 deep well plates. Sub-pools with a maximum of 96 ORFs each were prepared, and DNA was extracted using the QIAprep Spin Miniprep Kit (Qiagen). Each sub-pool was cloned into pDEST (pAWH or pBWH) destination vector by Gateway technology with LR clonase II enzyme mix (Thermo Fisher Scientific). Equimolar amounts of each pDEST-ORF sub-pool were mixed to generate the final pDEST H1001 library. Yeast transformations were performed as described (Yang et al., 2018). To minimize bias and cover the library size of 1001×1001, 48 yeast transformation were performed. Briefly, each pDEST-ORF was linearized with I-CeuI and I-SceI (New England Biolabs) at 37 °C for 16 h, followed by 20 min at 65 °C. For each transformation, 20 fmol of linear pAWH-ORF pool and 20 fmol of linear pBWH-ORF pool were co-transformed into freshly prepared competent Y2HGold cells according Clontech’s protocol. All transformations were pooled together and split into 2× 250 ml of RS and 2× 250 ml of RIS liquid-gel media containing 0.5% (w/v) Seaprep Agarose (Cultek), Minimal SD base (Clontech) and the indicated Amino Acid Dropout mix (Clontech). Cultures were grown in 5 L flasks for 60 h. Cells were harvested by centrifugation at 1600 g for 10 min, and washed once with PBS by spinning them down at 700 g for 5 min.

Library preparation for paired-end sequencing was performed as described in Yang et al. For each selection media, recombined yeast plasmid DNA, or pFAB (plasmid fusion of activation domain and binding domain), was extracted with 12 columns, with 4×10^7^ cells on each column, of Zymoprep™ Yeast Plasmid Miniprep II (Zymo Research). Yeast plasmid DNA was heated at 65 °C for 20 min. DNA from the 12 columns was pulled together and sheared by Covaris ultrasonication to a size of 1500 bp. DNA was end-repaired with NEBNext^®^ End Repair Module (New England Biolabs), purified with MinElute PCR Purification Kit (Qiagen) and circularized by an intramolecular ligation reaction using the Quick Ligation™ Kit (New England Biolabs)

To specifically amplify circular fragments containing 3′ ends of both a bait-ORF and a prey-ORF, and to add R1 and R2 Illumina adaptors, circular DNA was split in 20 PCR independent PCR reactions, with Q5 High-Fidelity 2X Master Mix (New England Biolabs), and primers Pr4seq_F_TS_R1 and Pr4seq_R_TS_R2 (synthesized by Integrated DNA Technologies, see Yang et al for primer sequence). R1/R2 PCR fragments were purified with Agencourt AMPure XP beads (Beckman Coulter). R1/R2 purified fragments were split in 12× 25 μl-reaction PCR reactions with Q5 High-Fidelity 2X Master Mix (New England Biolabs), and NEBNext^®^ Multiplex Oligos for Illumina^®^, Index Primers Set 1 (New England Biolabs). P5/P7 PCR fragments were purified with Agencourt AMPure XP beads (Beckman Coulter). Multiple independent PCRs were performed to minimize PCR bias. All steps of library preparation for paired-end sequencing were performed within the same day with freshly harvested yeast, to minimize yeast plasmid degradation and maximize the quality of the final product. The H47-library was screened 5 times and the RBPome library 11 times to ensure a maximized sampling complexity.

### rec-Y2H data analysis

Raw paired-end reads (read 1 and read 2) from RS and RIS conditions were analyzed with the rec-Y2H program from GitHub (https://github.com/lionking0000/recYnH). As described in (Yang et al., 2018), read 1 and read 2 were filtered, trimmed, and finally mapped with reference protein sequences with blanstn program. We calculated rec-Y2H interaction scores for H47 set and the large set as we previously described (avgIS). In order to increase the detection sensitivity (Figure S1X), we merged all the raw paired-end reads as one single experiments before applying noise filter and reconstructing null matrix. Then the noise-filtered signal is normalized by the null matrix to generate the interaction score (sumIS) matrix. To calculate the IS differences for the RBM10, SRRM3, and SRRM4 mutation data. We firstly calculated interaction scores of each case and then subtract the interaction score of wild type proteins for each corresponding binding partner. The interaction enrichment matrix was calculated by the observed number of interactions between proteins that belong to two classes divided by the number of interactions from the randomly shuffled same number of protein pairs. We used the same strategy to show the interaction enrichment between the same localized proteins, we calculated the null expected interaction from the randomly shuffled pairs.

### NanoBRET validation

pHTNW (Addgene #136403) and pNLF1W (Addgene #136404) have been previously described (Yang et al. 2018). ORFs of interest were cloned into pENTR vector by Gateway BP reaction and then transferred either to pHTNW or pNLF1W by Gateway LR reaction (Thermo Fisher Scientific) following the manufacturer’s protocol. LR reaction were transformed into 10 ul of OmniMAX™2 competent cells (Thermo Fisher Scientific). Positive clones were confirmed by restriction analysis using BsrGI enzyme and by sequencing.

HEK293T cells were plated in 24-well plates at a density of 1.4 × 10^5^ cells per well. Cells were transfected with 500 ng of pHTNW-ORF, 5 ng of pNLF1W-ORF, 0.75 μl of Lipofectamine 3000 and 1 μl of P3000 Reagent (Thermo Fisher Scientific). After 20 h, each transformation was re-plated in four wells of 96-well plates at a density of 1 × 104 cells per well for duplicate control and experimental samples (technical replicates), and PPIs were analyzed with NanoBRET™ Nano-Glo® Detection System kit (Promega) following manufacturer’s instructions. Each transformation experiment was performed at least twice. The corrected NanoBRET ratio was calculated according to the manufacturer’s instructions.

### Protein–RNA interaction data analysis

Interaction data for 94 RNA-binding proteins for which eCLIP data were available and which were assayed in our screen were obtained from the ENCODE Project’s data portal in narrowPeak BED format (Van Nostrand et al., 2016; Sloan et al., 2016), see Supplementary Table S3 for sample accessions. We used the “reproducible” set of interactions (Van Nostrand et al., 2020), where a peak must be found with precisely the same start and end coordinates in both replicates in a given cell type to be included. Interaction data were filtered using thresholds of p<10^−3^ and ≥8-fold enrichment over “SMInput” (paired size-matched input) (Van Nostrand et al., 2020). The genomic coordinates of the eCLIP peaks described above were mapped to genes they overlapped with from Ensembl release 91 (Zerbino et al., 2018). All plots were generated using version 1.3.0 of the tidyverse packages (principally ggplot2 3.3.2) in R 3.6.0 (see https://github.com/langbnj/rbpome for scripts). Other packages used were reshape2 1.4.4, glue 1.4.1, broom 0.5.6, scales 1.1.1, Hmisc 4.4-0, and corrplot 0.84.

Positive controls were defined as RBP pairs which were identified as direct interactors by at least two studies in BioGRID release 3.5.185 (May 2020) (Oughtred et al., 2019). For this, studies using co-crystal structures, reconstituted complexes or two-hybrid approaches were considered capable of reporting direct interactions. The resulting controls were FMR1-FXR2, HNRNPK-QKI, RBFOX2-QKI, SFPQ-NONO, and U2AF1-U2AF2.

Random RBP pairs were used as a negative control. For this, we randomly chose two proteins out of the 94 RBPs with available ENCODE eCLIP data to arrive at a unique set equal in size to the interactions identified in our screen (resampling without replacement). We performed 100 random resamples to arrive at resampling p-values <0.01.

### RBP target set similarity quantification

To quantify the similarity of target sets between pairs of RNA-binding proteins (Figure 2B & 2F), we employed the Jaccard index (defined as the size of the intersection of the two sets, i.e. RNA target genes bound by both RBPs, divided by the size of their union, i.e. the total number of RNA target genes bound by at least one of the RBPs). To reduce redundancy and a potential bias towards genes encoding multiple alternative transcripts through alternative splicing or initiation sites, this analysis was performed at the gene level, as determined by peak overlap as described above. We compared the Jaccard index values of the RBP pairs whose interactions were identified in our screen to random pairs (negative control as described above). We used both a resampling p-value across 100 resamples and a one-tailed Wilcoxon rank-sum test between the screen hits and the pooled resamples to ascertain whether the observed RBP pairs displayed higher target set similarity than random pairs. To ensure comparability of the box plots, only one of these random resamples is shown in Figure 2B.

### Conditional probability of co-binding

The conditional probability of co-binding was defined as the probability p(A|B) of protein A binding a transcript from a gene of interest, given protein B binding a transcript from the same gene (Figure 2C & 2F). As for the target set similarity above, this analysis was performed at the gene level to reduce bias arising from the number of alternative transcripts per gene. The statistical analysis was likewise performed as above.

### RBP binding distance measurement

To determine the distance observed between the binding sites of a pair of RNA-binding proteins, we compared the positions of the 5’ ends of all their peaks within a given gene and used the overall per-gene minimum as a metric of proximal binding. The 5’ end of an eCLIP peak is predicted to correspond to the site of the covalent cross-link between an RBP and RNA (Dominguez et al., 2018) and should therefore be the closest approximation of the actual interface. The cumulative density functions (CDFs) for the positive control (described above), identified interactors, and for 100 sets of random RBP pairs (negative control, described above) were plotted using the stat_ecdf function of the ggplot2 R package (Figure 2D). Additionally, we generated control binding distance datasets for each set by randomizing the positions of the peaks within each target gene (i.e. maintaining both peak number and gene lengths).

### Calibration of proximity binding threshold using positive controls and random binding data

In order to determine a threshold below which an RBP pair binds in extreme enough proximity to indicate binding together as a complex, and to distinguish this from incidental nearby binding, we used the five positive control RBP pairs listed above as a gold standard. For these positive controls, which are known to bind their RNA targets as complexes, we determined the threshold on their cumulative density function (CDF) where the difference between their binding site distances and those observed among the random pairs was maximized (≤54 nt, Figure 2E). For each set (positive control, identified interactions and random RBP pairs), this binding site distance difference (Δ) was calculated by subtracting the binding site distance CDF for the randomized positions (described above). In this analysis, distances above 1000 nt were discarded as these would require long-range RNA structures in order to be bound by an RBP complex, in which case primary sequence distance would not be predictive of binding as a complex.

### Test for significant adjacent binding

To determine whether a given pair of RNA-binding proteins displayed closer binding than expected by chance, we generated violin plots showing the binding distances of a given RBP in a pair relative to those of another (e.g. “FXR2 relative to FMR1”) (Figure 2G–I). For each binding site of the reference RBP (e.g. FMR1) within a gene, we determined the distance to the closest peak of the other (e.g. FXR2). We generated a control dataset by randomizing the positions of the proteins’ binding sites within each target gene (i.e. maintaining both binding site number and gene length distribution). The “close proximity” binding threshold of 54 nt is highlighted in the figures in order to visualize the fraction of binding sites which may be bound by the pair as a complex. Two statistical tests were used in concert to determine whether the number of binding sites in close proximity (≤54 nt) observed for a given pair (and pair orientation) were significantly higher than for randomized peak positions: a simple resampling-based test (100 resamples), and a variant resampling-based test which used a one-tailed Wilcoxon rank-sum test to compare each resample to the observed data (p<0.05) and used the number of successful tests to arrive at the equivalent of a “resampling p-value”. This second test proved more stringent than the first and tended to reject problematic pairs that had very few binding sites overall, and whose low fraction of binding sites in immediate proximity the first test had nonetheless considered significant. Requiring both tests to be successful filtered out all such problematic pairs. We decided to use this methodology rather than discarding pairs without a minimum number of binding sites in immediate proximity, which would have achieved a similar effect only at >670 sites.

### Network of RBPs binding in close proximity and potentially as complexes

A network showing genes with eCLIP protein–RNA interaction data from the ENCODE Project was generated using Cytoscape 3.8.0 (Figure 3A). This network incorporated only those pairs which bound in close proximity (≤54 nt) more frequently than expected from randomized peak positions, as determined using both tests described above for at least one pair orientation (i.e. either RBP A binding near RBP B, or RBP B binding near RBP A). Known complexes with direct interaction evidence were obtained from BioGRID 3.5.185. As above, studies using co-crystal structures, reconstituted complexes or two-hybrid approaches were considered capable of reporting direct interactions. Subcellular localization information was obtained from the Human Protein Atlas (Thul et al., 2017), dataset dated December 17^th^, 2019).

### Conversion of genomic binding coordinates to meta-mRNA positions

Meta-mRNA plots were generated by using transcript annotation from Ensembl 91 (Zerbino et al., 2018). To reduce bias stemming from the number of known alternative transcripts per gene, which can vary widely, we retained only the most highly expressed protein-coding RNA transcript per Ensembl gene. This transcript was chosen using TPM (transcripts per million) expression data collected as part of the ENCODE Project (accessions ENCSR000CPE and ENCSR000CPH for HepG2 and K562 cells, respectively) (Sloan et al., 2016). Each eCLIP peak falling within this most highly expressed transcript for a given gene was assigned to an mRNA region using the 5’ end of the peak (see above for the rationale: the 5’ end should be most representative of the interface). The regions used were: 5’ UTR, first exon, first intron, internal exons, internal introns, final exon, final intron, and 3’ UTR. Each peak’s 5’ position was scaled to a range of [0, 1] within its assigned region, allowing interpretation across transcripts and genes of differing lengths.

### Generation of meta-mRNA probability density profiles

Probability density profiles were generated using ggplot2. To determine y-axis error ranges, we performed 1000-fold resampling with replacement on the target genes in each plot and plotted the 95% confidence interval around the median (Figure 3C, E, and G). In the probability density plots, the within-region x-axis scaling mentioned above was changed to a range of [0.1, 0.9] to more clearly demarcate the boundary between regions, and to allow clear interpretation of whether a probability density peak lies at the 3’ end of one region or at the 5’ end of another.

### Meta-mRNA correlation heat maps

Meta-mRNA probability density coordinates were obtained using the R density function at a bandwidth of 1/320. Correlation heat maps were then generated from Pearson correlation coefficients via the R corrplot package (as mentioned above), with hierarchical clustering use as the ordering function.

### Identifying known target sequence motifs for RBPs

We predicted known human RNA-binding protein motifs within eCLIP peak regions using the FIMO program (Grant et al., 2011) from the MEME Suite (Bailey et al., 2009). Peak regions were extended 50 nt upstream from their 5’ end, following the reasoning of Dominguez *et al.* that the 5’ end of the eCLIP peak represents the UV cross-linking site between protein and RNA, implying that the actual binding interface can be upstream of it (Dominguez et al., 2018). A comprehensive set of literature-derived motif position weight matrices (PWMs) were obtained from the ATtRACT database (Giudice et al., 2016), Dominguez *et al.*’s RNA Bind-N-Seq (RBNS) study (Dominguez et al., 2018), the RNAcompete study (Ray et al., 2013), and the CISBP-RNA (Ray et al., 2013), RBPDB (Cook et al., 2011)and RBPmap (Paz et al., 2014)databases. For RNAcompete and RBPmap, we used the PWMs included with the MEME suite. Motif PWMs from all other sources were reformatted to MEME’s format using custom Perl scripts, so they could be used by FIMO. Known motifs were available for a total of 29 RBPs (74 interacting pairs). For FIMO, a uniform background sequence model was used (equiprobable A, C, G, U) since inferring a background model for the peak regions, either for all or for individual RBPs, resulted in very low motif hit rates, apparently since motifs are often repeated within peak regions. We used FIMO with a p-value threshold of 0.001. We did not use q-values due to the variation in the number of binding sites between different RBPs, which is also evident between eCLIP biological replicates and therefore appears random. We considered it unreasonable to penalize eCLIP experiments that may have achieved higher sensitivity and reported larger numbers of binding regions, and considered FIMO’s p-values to be more comparable between experiments than q-values.

### Determining motif presence and count correlations

To determine whether known motif presence and absence can be used as an additional quality measure for eCLIP binding sites where both RBPs bind in close proximity (≤54 nt), we tested whether motif presence for RBP A in its binding region correlates with the presence of a known motif for RBP B using Fisher’s exact test. Similarly, since there is frequently more than one occurrence of a given motif in a binding region, we tested whether the number of predicted motifs is linearly correlated. For this, we used Wilcoxon rank-sum tests (i.e. Spearman) and Pearson product-moment correlation tests. As visible in Figure 4B, all significant correlations are positive.

### Determining whether RBP sequence motif usage differs at co-bound sites

To determine whether RBPs bind different sequence motifs depending on their likely complex partners, we tested all possible combinations of motifs for a given pair of RBPs. For each RBP pair A-B and motif pair a-b, we identified those binding sites where A and B bind within 54 nt (as determined by the 5’ ends of their eCLIP peaks, and as described above). We then counted the number of binding sites of RBP A that contained motif a, where RBP B’s closest binding site contained motif b. As a control, this number was compared to a random sampling and pairing of RBP A’s and RBP B’s binding sites, from target genes where only RBP A or RBP B binds. We sampled the same number of control sites as there were observed sites (with replacement). Both numbers were converted to pseudocounts by adding 1, and an enrichment ratio was calculated by dividing the observed number of occurrences by the control. We thus calculated a pseudocount enrichment ratio of motif pair a-b’s occurrence at sites where RBPs A and B bind in proximity (≤54 nt), and likely as a complex, compared to the background probability of observing motif pair a-b. To obtain a confidence range for this motif occurrence ratio, as shown in each box plot in Figure 4C, we then performed a 100-fold resampling (with replacement) across both the observed and control sets. Motif logos were generated from the PWMs using meme2images from the MEME Suite.

### Interaction network analysis for RNA metabolic processes

To generate the interaction networks sorted by RNA metabolic step, screen hits above a cut-off of 7.1 were intersected with lists of proteins belonging to different metabolic steps. All protein lists were obtained from the Reactome resource, except for stress granule and processing body proteins (RNA granule database) and cytoplasmic mRNA transport (literature curated list). For the stress granule network, the sumIS threshold was raised to 10 as otherwise the network would have been too complex for graphical representation. Networks were generated using Cytoscape V3.7.2 and the yFiles hierarchic layout addon.

## Supplementary Materials

Figure S1. Additional screen benchmarking. Related to Figure 1.

Figure S2. Intersection of eCLIP and RBP PPI data. Related to Figure 2.

Figure S3. Correlation of meta pre-mRNA profiles. Related to Figure 3.

Figure S4. Extended RBP-pair di-motif preference. Related to Figure 4.

Figure S5. Additional RBP networks. Related to Figure 5.

Figure S6. Reproducibility of RBP-mutant interactions. Related to Figure 6.

Table S1. Screen input library. Related to Figure1.

Table S2. Screen results. Related to Figure 1.

Table S3. eCLIP sample metadata. Related to Figure 2.

Table S4. Random resamples NanoBRET data. Related to Figure 2.

Table S5. RBP co-binding statistics. Related to Figure 2 and 3.

Table S6. Identified interactions sorted by process. Related to Figure 5.

Table S7. RBP mutants screen results. Related to Figure 6.

Supplementary Data1. Cytoscape file containing all rec-Y2H and NanoBRET data.

