## Supplemental figures and legends for "rec-Y2H matrix screening reveals a vast potential for direct protein-protein interactions among RNA binding proteins"

**This PDF file includes:**

Figures S1 to S6 with legends

Captions for supplementary Cytoscape data file 1

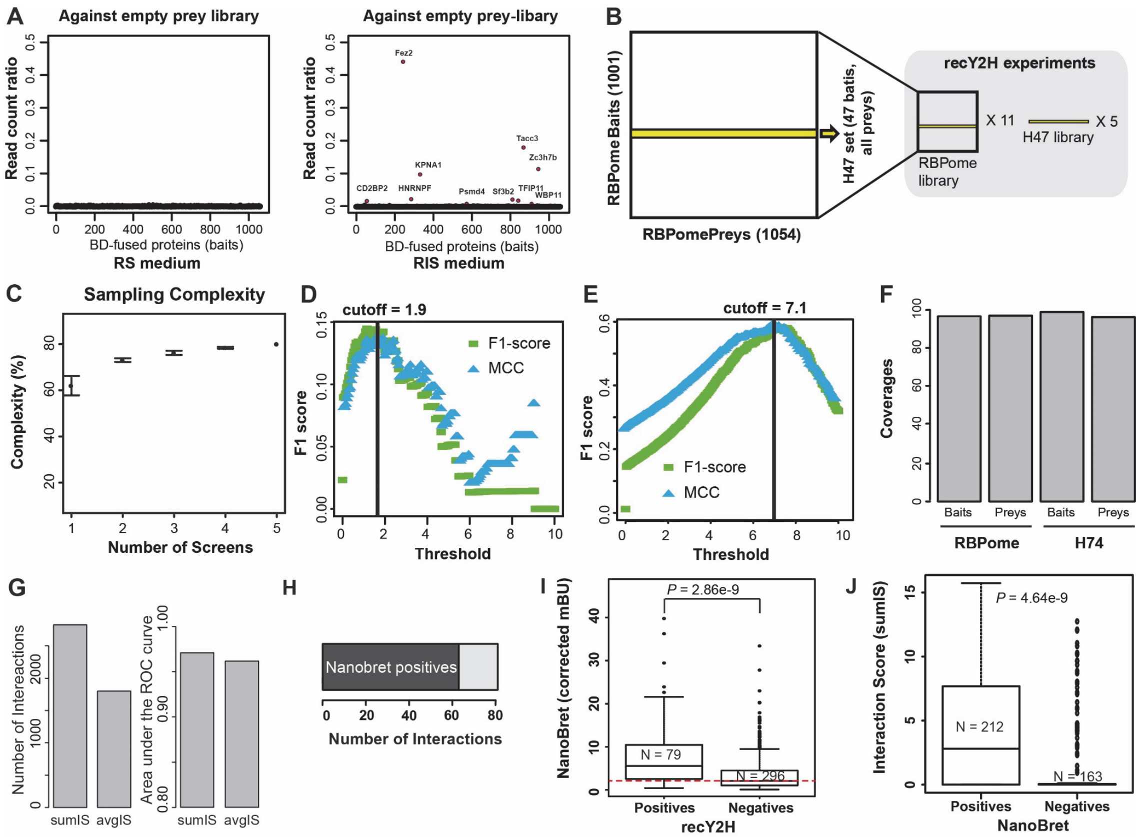


**FigureS1: Additional screen benchmarking. (A)** A priori detection of auto-activators. The entire bait-library was screened against an empty prey-vector. The RS (recombination–selection) conditions showed no significant bias of reads detected per bait. The RIS (recombination–interaction–selection) condition showed that some baits produced reads despite of the absence of prey-interaction partners and are hence classified as auto-activators and removed from the bait library for all further screens. **(B)** Scheme of the libraries screened. The RBPome-library consists of 1001 Baits (after the removal of auto-activators) and 1054 Preys. For the small-scale rec-Y2H screens, we used a library of 47 Baits and 1054 Preys, called H47-library. **(C)** Sampling complexity as a function of the number of H47 screens. Sampling complexity is the fraction of pairs that form out of all possibilities under RS conditions. **(D)** The optimal interaction score cut-off for the H47-library was determined based on the harmonic average of precision and sensitivity (F1-Score, green) and Matthews's Correlation Coefficient (MCC, cyan). We combined BioGRID and HIPPIE PPI databases to evaluate the performances. All pairs not present in any databases are defined as none-interacting pairs. **(E)** The optimal interaction score cut-off for the RBPome-library screen was determined based on the maximal F1 score indicating the performance of distinguishing the H47 set positives and negatives. **(F)** Detection of baits and preys in the RBPome- and H47-screens. **(G)** Benchmarking of the sumIS and avgIS based on the number of detected interactions and performance. For sumIS, 2825 interactions were above the cutoff (=7.1) for the whole RBPome screen. For avgIS 1799 interactions were above the cutoff (=1.7) for the same RBPome screen. We note that the interactions are distinguishing the orientations (Px-AD—Py-BD or vice versa) and the score thresholds were determined based on the MCC values with H47 results. Both of the scores show high performance based on H47 set as golden positive and negative set, 0.9709 for sumIS and 0.9614 for avgIS, respectively. **(H)** Benchmarking of the RBPome screen with NanoBRET validations. 375 interactions were tested. Interactions scored positive in the rec-Y2H RBPome screen had a significantly higher average NanoBRET score than pairs tested negative in rec-Y2H. **(I)** NanoBRET scored positives had a significantly higher average Interaction score than pairs in the rec-Y2H RBPome screen **(J)** The fraction of NanoBRET-validated interactions above cut-off (sumIS=7.1) **(K)** A network of NanoBRET-validated interactions with a sumIS score above cut-off.

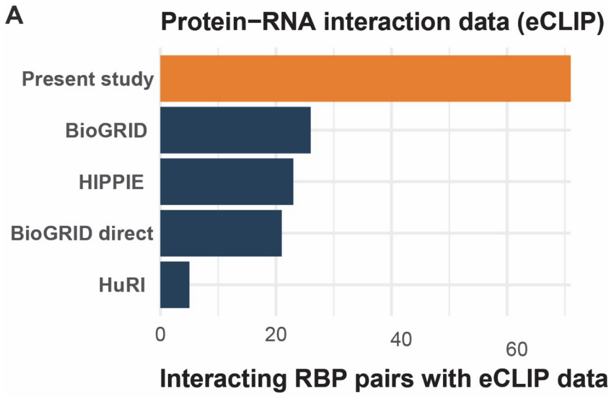


**Figure S2: Intersection of eCLIP and RBP PPI data. (A)** The number of protein-protein interactions among RBPs for which eCLIP data exists.

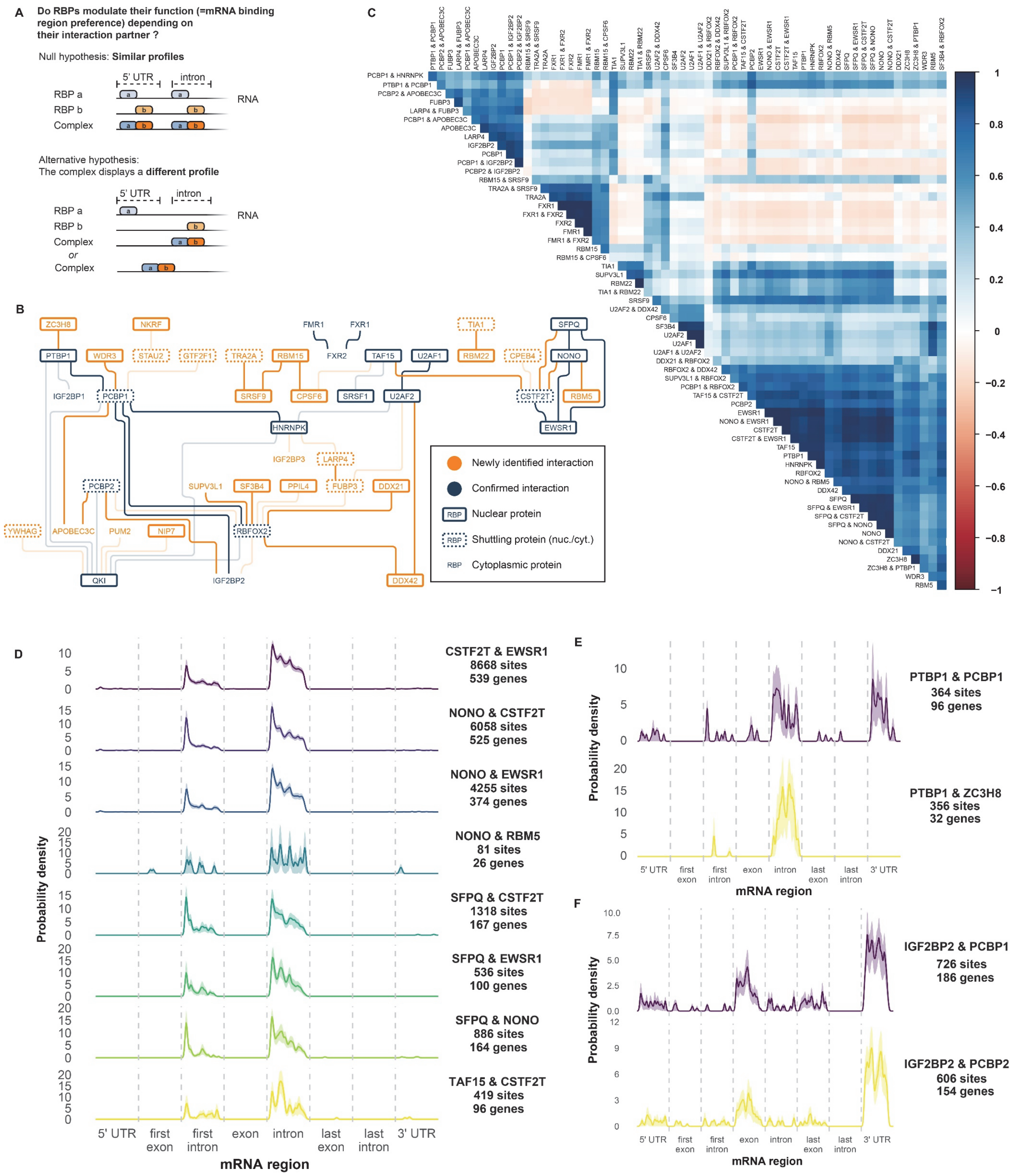


**Figure S3: meta pre-mRNA profiles. (A)** Illustration of the hypothesis tested in Figure 3. **(B)** Network graph of RNA-binding proteins with eCLIP protein–RNA interaction data. Connections indicate statistical support for binding in proximity: transparent lines, weak support by at least one type of test in one pair orientation, opaque lines: strong support by both types of test used in at least one pair orientation. Subcellular localisation is indicated by gene symbols framed either by a continuous border (nuclear proteins), a dotted border (shuttling, i.e. sometimes nuclear), and no border (always cytoplasmic). Interactions newly identified in our screen, and the proteins newly joined by them to the network, are highlighted in orange. **(C)** Correlation matrix of PDF pre-mRNA profiles from all induvial RBPs and RBP pairs with significant proximity binding within 54 nt (Figure 3A). **(D)** PDF plots showing meta-pre-mRNA binding profiles of the NONO-cluster detected in our screen. All proteins, which have multiple interactions amongst each other (Figure S3B), show a highly similar binding profile. **(E)** PDF plots showing the meta-pre-mRNA biding of PTBP1 depending on its interactions partner. **(F)** PDF plots showing the meta-pre-mRNA biding the close homologous PCBP1 and PCBP2 with the same interaction partner.

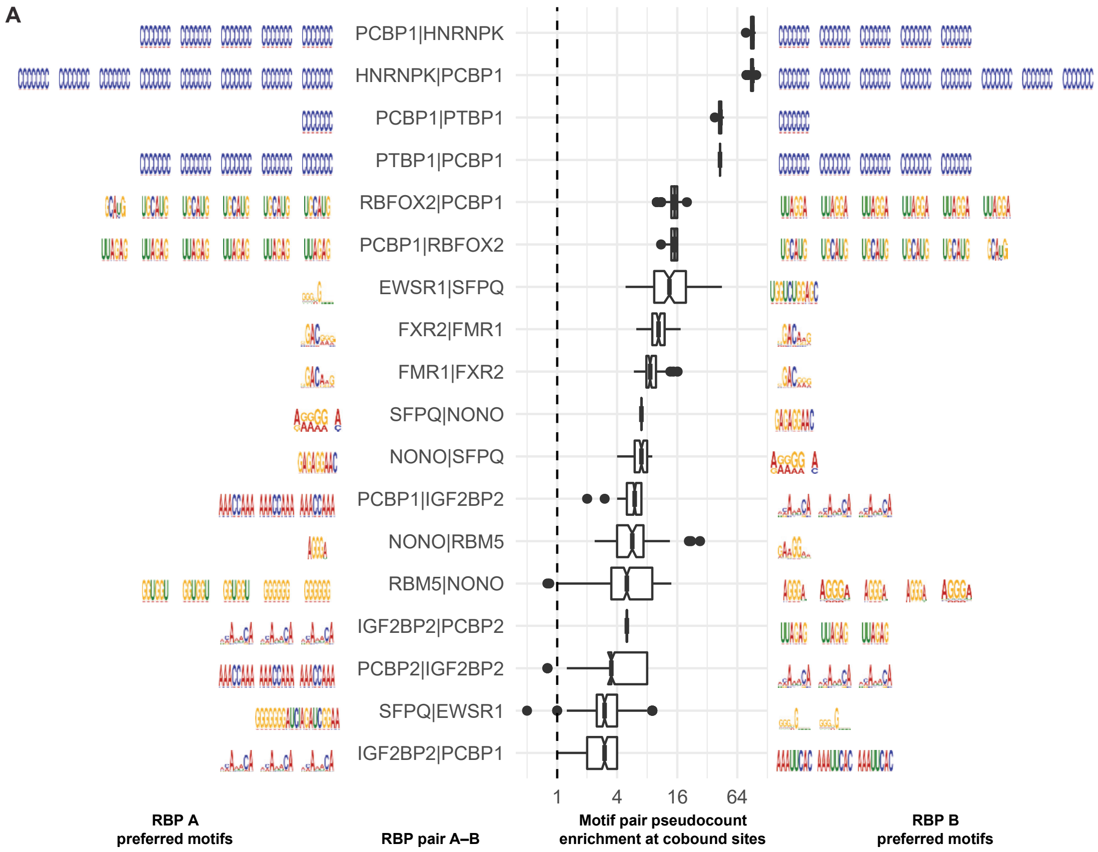


**Figure S4. Extended RBP-pair di-motif preferences. (A)** For each RBP pair, the preferred combination of motifs for RBP A (left) and RBP B (right) when both bind in proximity is shown as a sequence logo. The box plots show the pseudocount ratio indicating how frequently a given motif combination occurs when two RBPs bind in close proximity as compared to an independent-binding background derived from target genes where only one of the RBPs binds. Where median ratio ties are present (multiple motif combinations with the same median ratio as indicated on the box plot), all top-ratio motifs are shown.

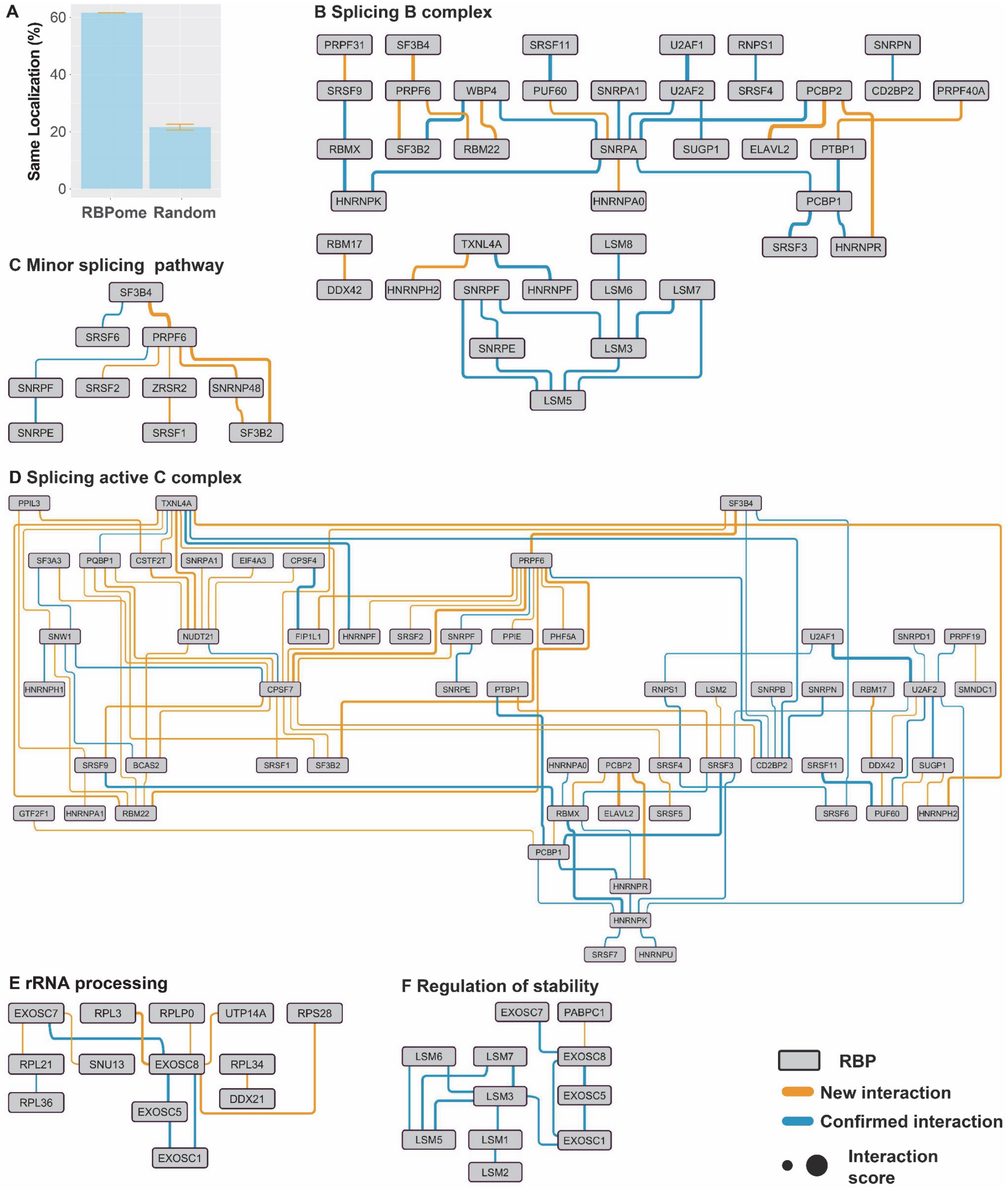


**Figure S5: Additional RBP networks. (A)** The tendency of interactions among proteins with the same cellular localization. 60% of interactions found were among the same localization proteins. The randomly selected the same numbers of interaction pairs were tested as control. **(B-F)** Interactions detected by the RBPome screen, filtered by proteins involved in different RNA metabolic steps which were defined based on the reactome database.

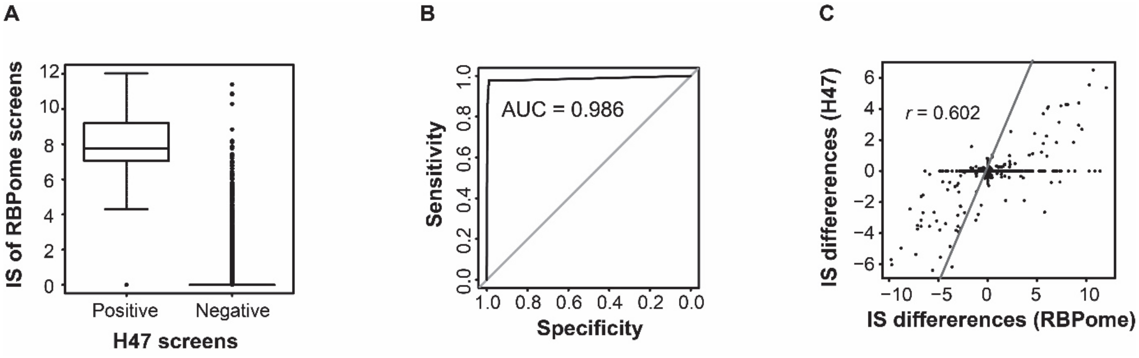


**Figure S6: Reproducibility of RBP-mutant interactions.** Benchmarking of reproducibility of interaction score changes in response to RBP mutation. **(A)** The interactions detected above F1-score based cut-off are positives and the others are negatives in the H47 set. Box plots show the distribution of sumIS from the full RBPome library screening for corresponding positive and negative interactions found in the H47 set. **(B)** Benchmarking the RBP mutant screen of the full RBPome library screens against the H47 calibration screen. The high score of Area Under the Curve (AUC) shows that screening two different libraries produce a highly similar result. **(C)** The interaction scores changes correlate in response to mutations between the full RBPome library screen and H47-calibraration library screen.

**Supplementary Data1.**

Cytoscape file containing all rec-Y2H and NanoBRET data. We provide two different RBP interaction networks based on sumIS threshold scores (7.1 and 4.5). Novel interactions detected by rec-Y2H are indicated by orange lines and known interactions by blue lines. NanoBRET validated interactions are indicated as double lines. Node color indicates protein location. Nuclear localized proteins, cytoplasmic proteins, nuclear and cytoplasmic dual-localized proteins are distinguished by red, orange and pink. The width of the lines connecting proteins indicates sumIS scores from 4.5 to 17.2.
